# Novel transcription factor BTNL9 enhances tumor suppression and drug sensitivity in non-small cell lung cancer through cell cycle regulation

**DOI:** 10.1101/2025.03.02.640945

**Authors:** Wooi Loon Ng, Pedram Yadollahi, Hwa Jin Cho, Mi Seon Kang, Inhak Choi

## Abstract

**Background:** Butyrophilins (BTNs) are immunoglobulin superfamily proteins involved in immune regulation. Among them, BTNL9 has unique structural features, including a bZIP-like domain, suggesting a potential transcriptional role. While BTNL9 is known to suppress T cell activation, its function in cancer remains largely unexplored. Recent studies suggest it may inhibit tumor progression and correlate with improved prognosis in multiple cancers. However, its molecular mechanisms and regulatory impact on lung cancer remain unclear. This study investigates the role of BTNL9 as a transcription factor and its implications for tumor progression and therapy.

**Methods:** ChIP-seq identified BTNL9-binding sites, followed by RNA-seq to assess transcriptomic profiles and validated by western blot. Drug sensitivity was evaluated through cytotoxicity assays. A xenograft model was applied to assess the effect of BTNL9 on tumor growth. TCGA data analysis examined correlations with survival, cell cycle regulators, and immune infiltration.

**Results:** ChIP-seq identified 26,610 BTNL9 binding peaks, mapping to 9,707 genes near transcription start sites. RNA-seq and western blotting showed BTNL9 regulates cell cycle (*E2F1*, *CDKN1A*, *CDK1*, *CDC25C*, *FOXM1*), DNA replication (*MCM2*/*3*/*7*, *ORC6*), and p53-related transcription (*BBC3*, *GADD45A*). Integrative analysis found that 74.8% of differentially expressed genes were directly regulated by BTNL9. Functionally, BTNL9 overexpression induced cell cycle arrest, reduced proliferation, and suppressed tumor growth in vivo. BTNL9 enhanced bortezomib sensitivity in both A549 and NCI-H460 cells, with etoposide effects being more pronounced in A549. Higher BTNL9 levels strongly suppressed the expression of FOXM1, CDC25C, CDK1, CDK2, CCNA2 and CCNB1 and negatively correlated with these markers in LUAD TCGA data. Elevated BTNL9 expression was associated with improved survival, complete remission, and increased immune infiltration, including macrophages, CD8+ T cells, NK cells, and B cells in cancer tissues.

**Conclusions:** BTNL9 functions as a transcription factor, suppresses tumor growth, and enhances drug sensitivity. Its correlation with survival and immune infiltration suggests potential role as a tumor suppressor and predictive biomarker for chemotherapy response.

## INTRODUCTION

Butyrophilins (BTNs) constitute a group of transmembrane proteins that belong to the immunoglobulin superfamily and are structurally homologous with immune-regulatory B7 family proteins. BTNs participate in modulating immune responses via costimulatory or inhibitory signals and have been linked to autoimmune inflammatory diseases and lipid metabolism (1–5). The human genome harbors 13 BTN/BTN-like (BTNL) genes (6), among which BTNL9 has attracted attention due to its unique features. BTNL9 has been demonstrated to function as a negative regulator of T cell activation in Jurkat T cells and murine models (7, 8). The IgV1 domain of BTNL9 has the capacity to interact with and bind to CD8 T cells, B cells, and NK cells, thereby suppressing immune modulation (9). Most BTN/BTN-like genes possess distinct IgV, IgC, and SPRY/B30.2 domains. While BTNL9 presents with one IgV domain, it lacks a clear IgC domain, unlike the highly structurally similar BTNL3 and BTNL8. Moreover, the absence of two critical cysteine residues at positions 173 and 227, which are crucial for the stabilization of the IgC domain through disulfide bonding, indicates that BTNL9 might not have a typical immunoglobulin structure (6). Another unique feature of the protein sequence of BTNL9 is that it exhibits a basic leucine zipper domain (bZIP domain), a common feature among transcription factors, but has just a single leucine heptad repeat that encompasses a coiled-coil region (current data in this study). The BTNL9 gene is located at the 5q35.3 locus, which is positioned near the terminus of the q arm of chromosome 5, an area known for its high density of transcription factor (TF) genes (10). This suggests that BTNL9 might function as a transcription factor. In addition to these unique structural features of BTNL9, recent in vitro studies have revealed that it has inhibitory effects on cell proliferation, invasion, and migration in melanoma and breast cancer cell lines (11, 12). Further, transcriptomics data mining has suggested a link between higher BTNL9 expression and good prognosis of lung, breast, bone, and colon cancer, as well as indicated its potential role in tumor suppression (13–18).

In this study, we present, for the first time, compelling evidence that BTNL9 acts as a novel transcription factor in lung cancer. It includes a global mapping of BTNL9’s binding sites and its active involvement in regulating genes critical for DNA replication and cell cycle. Furthermore, BTNL9 appears to steer genes tied to p53-regulated transcription in a p53-dependent manner. Using xenograft model, we showed that BTNL9 inhibits tumor growth in vivo. We also demonstrated that BTNL9 protein level in cancer cells is associated with etoposide and bortezomib treatment sensitivity. Analysis of clinical data from lung cancer patients revealed that higher BTNL9 expression was associated with better survival and improved rates of complete remission in those undergoing primary treatment. Overall, our study offers a deeper understanding of transcriptional regulation and tumor suppression dynamics, hinting at a broader mechanism for cancer control.

## MATERIALS AND METHODS

### Ethical statement

The animal study is approved by the Institutional Animal Care and Use Committee of Inje University under the project title” Study of BTNL9 cancer suppression mechanism” (Institutional Review Board# 2023-024), and fully complied with the accepted standards for the care and use of laboratory animals.

### Cell culture

Human lung carcinoma cell lines A549 (RRID:CVCL_0023), NCI-H460 (RRID:CVCL_0459) and human embryonic kidney cell 293T (HEK-293T, RRID:CVCL_0063) were obtained from Korean Cell Line Bank (KCLB) and American Type Culture Collection (ATCC). A549 and NCI-H460 were cultured in RPMI-1640 medium (Cytiva, #SH30027), and HEK293T was cultured in Dulbecco’s Modified Eagle Medium (DMEM) supplemented with 10% fetal bovine serum (FBS) (Cytiva, #SH30919.03) and 1% penicillin/streptomycin (Cytiva, #SV30010). All the cell lines were maintained at 37 °C with 5% CO_2_and sub-cultured upon reaching approximately 80% confluency.

### Plasmid construction of human *BTNL9*

Optimized human BTNL9 cDNA (NM_152547.5) cloned in pcDNA3.1 (+) vector was purchased from Invitrogen and used as a template for further plasmid constructions. Amplification of human *BTNL9*, flanked by 5’ and 3’ end for the pLJM1-eGFP vector (Addgene, RRID:Addgene_19319), was performed via PCR using PrimeSTAR GXL DNA Polymerase (Takara, # R050A) and primers were listed in the Table S8. The vector system was slightly modified by removing the GFP, and either AM-tag or FLAG-tag was fused to the C-terminus of BTNL9. Plasmid was digested with *Nhe*I, *Pml*I, or *Eco*R1 and PCR products were cloned into the vector via Gibson assembly. Recombinant plasmids expressing BTNL9-AM-tag, BTNL9-FLAG-tag or without tag of BTNL9 were constructed.

### Lentivirus production and infection

Lentiviral particles were produced using 2nd generation packaging system according to previous published protocol with minor modification (19). Three days after transfection, viral supernatant was harvested, subsequently filtered through 0.45 μm filter to remove cell debris, and then mixed with Lenti-X concentrator at a 3:1 ratio (Takara, # 631231). After incubation overnight at 4 °C, the mix was centrifuged at 1,500 x g for 45 minutes at 4 °C. The concentrated lentivirus was re-suspended in appropriate volume and kept at – 80°C. A549 and NCI-H460 were subsequently transduced with lentiviruses carrying either an empty vector or overexpression plasmids (pCMV-BTNL9-AM-tag, pCMV-BTNL9-FLAG-tag, pCMV-BTNL9) in the presence of 8 ug/mL hexadimethrine bromide (Sigma-Aldrich, # H9268) for overnight. Fresh medium was replaced on the next day and cells were harvested as assigned.

### Chromatin Immunoprecipitation sequencing (ChIP-seq) and ChIP-qPCR

ChIP DNA was prepared using Tag-ChIP-IT (Active Motif, #53022) according to the manufacturer’s protocol. A549 and NCI-H460 cells were infected with lentivirus overexpressing pCMV-BTNL9-AM-tag and cross-linked with 1% formaldehyde fixative solution on Day 3 before quenched after 15 min. Following 2 rounds of washing, cell pellets were incubated in Chromatin Prep Buffer for 10 min, followed by 20 strokes of processing with a chilled Dounce homogenizer. After centrifugation, cells were resuspended in ChIP buffer before sonication to yield fragments between 200-500 bp in length. Antibodies such as anti-AM-tag (Active Motif, #91111, RRID:AB_2793779), anti-mouse H3K27ac (Thermo Fisher Scientific, #MA5-23516, RRID:AB_2608307) or mouse IgG2a isotype control (Thermo Fisher Scientific, # 02-6200, AB_2532943) were added to 30 µg sheared chromatin and incubated at 4 °C overnight. ChIP DNA and input DNA were extracted using the column provided by this kit. Both ChIP DNA and input DNA were either analyzed with ChIP-qPCR or sent to ChIP-seq service provider (BGI Genomics, Hong Kong) for further processing and analysis. ChIP-DNA was sequenced using the DNBSEQ™ MGISEQ-2000RS sequencing technology platforms. ChIP-DNA or input were treated to repair DNA end before 3’-dA overhang and ligation of sequencing bubble adaptor. After amplification, heat separation and single strand circular to generate library, finally DNA sample were proceeded for sequencing. Data filtering were proceeded including removing adaptor sequences, contamination and low-quality reads from raw reads. After the filtering step, the clean data was mapped to the reference genome GRCh37/hg19 using SOAPaligner/SOAP2 (Version: 2.21t, RRID:SCR_005503). Peak calling was analyzed by Model-based Analysis of ChIP-seq v1.4 (MACS, RRID:SCR_013291), with regions identified as peaks when they had a *p* < 1.0E-05. Significant peaks were defined with more stringent threshold of Benjamini-Hochberg FDR < 0.005. Calculation of ChIP-DNA enrichment was based on standard of percentage of input. 10% of sheared chromatin as input was purified together with final ChIP-DNA. Equal amount of ChIP-DNA from IgG2a isotype control and BTNL9-AM-tag pulled down groups were used as template in ChIP-qPCR. Primers were listed in Table S8.

### MEME-ChIP DNA motif analysis

Both BTNL9 ChIP peak replicate samples were further processed to meet the criteria before being submitted for motif analysis. Peaks with a Benjamini-Hochberg FDR < 0.005 were selected to extract 500 bp from the central region of each peak and generated final BED format. In the CentriMo option within the MEME-ChIP software interface (RRID:SCR_001783), a maximum region width of 200 bp was set for motif analysis. Only motif sequence appeared from both replicate samples analyses was selected as the conserved motif in this study.

### RNA-sequencing (RNA-seq)

Total RNA was isolated using Trizol reagent (Thermo Fisher Scientific, # 15596026) and sent to RNA-sequencing service provider (Ebiogen). RNA quality was assessed by Agilent TapeStation 4000 system (Agilent Technologies), and RNA quantification was performed using ND-2000 Spectrophotometer (Thermo Fisher Scientific). Construction of library was performed using QuantSeq 3’ mRNA-Seq Library Prep Kit (Lexogen) according to the manufacturer’s instructions. High-throughput sequencing was performed as single-end 75 sequencing using NextSeq 550 (Illumina). Sequencing raw data was analyzed by ROSALIND^®^ (https://rosalind.bio/, RRID:SCR_006233), with a HyperScale architecture developed by ROSALIND, Inc. (San Diego, CA). Reads were trimmed using cutadapt1. Quality scores were assessed using FastQC2. Reads were aligned to the Homo sapiens genome build hg19 using STAR3. Individual sample reads were quantified using HTseq4 and normalized via Relative Log Expression (RLE) using DESeq2 R library. Read Distribution percentages, violin plots, identity heatmaps, and sample Multidimensional scaling (MDS) plots were generated as part of the QC step using RSeQC6.

DEseq2 (RRID:SCR_015687) was also used to calculate fold changes and *p*-values and performed optional covariate correction. Clustering of genes for the final heatmap of differentially expressed genes (DEGs) was done using the PAM (Partitioning Around Medoids) method using the fpc R library. Hypergeometric distribution was used to analyze the enrichment of pathways, gene ontology, domain structure, and other ontologies. The topGO R library was used to determine local similarities and dependencies between GO terms in order to perform Elim pruning correction. Several database sources were referenced for enrichment analysis, including Interpro9, NCBI10, MSigDB11,12, REACTOME13, WikiPathways. Enrichment was calculated relative to a set of background genes pertinent to the study.

### RNA extraction and cDNA synthesis

Total RNA for qPCR was extracted using TaKaRa MiniBEST Universal RNA Extraction Kit (Takara Bio, #9767A) according to the manufacturer’s protocol. RNA concentration and quality were measured with Nanodrop 2000 (Thermo Fisher Scientific). cDNA was synthesized from 2 μg of total RNA in 20 μl reactions using High-Capacity cDNA Reverse Transcription Kit (Thermo Fisher Scientific, #4368814). After synthesis, the cDNA was diluted 4 times with double distilled water and stored at −20 °C.

### Quantitative real-time PCR (RT-qPCR)

Real-time PCR was performed in a CFX Opus 96 Real-Time PCR System (Bio-Rad) using KAPA SYBR FAST Universal Kit (Sigma-Aldrich, #KK4601). For quantification using comparative CT method of gene expression, corresponding primers used in this study are listed in Table S8. All reactions were performed in triplicate and normalized to GAPDH, which served as a control housekeeping gene.

### Immunofluorescence staining and confocal microscopy analysis

Both A549 and NCI-H460 cells were seeded on poly-l-lysine (Sigma-Aldrich, #P8920) coated slide overnight prior to the process for immunofluorescence staining. Cells were fixed with 4% Paraformaldehyde at room temperature for 15 min and permeabilized with 0.3% Triton-X-100 in 5% BSA for 15 min, followed by 5% goat serum for 1 h. Either 1:250 dilution of anti-rabbit BTNL9 antibody (LS-Bio, #LS-C399149) or rabbit IgG control was added for overnight incubation. Slides were washed and incubated at room temperature with 1:1000 diluted anti-rabbit IgG Alexa Fluor® 594-conjugated (Thermo-Fisher Scientific, # A32740, RRID:AB_2762824) for 1 h, followed by DAPI to counterstain nuclei. Slides were mounted with VECTASHIELD^®^ Antifade Mounting Medium (Vector Laboratories, #H-1000-10). All confocal images were captured with Nikon Confocal laser microscope A1+ at 40X lens.

### Western blot

Cells were lysed in RIPA Lysis and Extraction Buffer (Thermo Fisher Scientific, #89900) with Halt™ Protease and Phosphatase Inhibitor Cocktail (Thermo Fisher Scientific, #78440). The concentration of protein lysate was measured using Pierce™ BCA Protein Assay Kits (Thermo Fisher Scientific, #23225). 30 µg of protein lysate was resolved by 10% SDS-PAGE and transferred to Immobilon-PSQ PVDF Membrane (Sigma-Aldrich, #ISEQ00010). Transferred membranes were incubated with a 5% Blotting-Grade Blocker (Bio-Rad, #1706404), followed by 1:1000 dilution of primary antibodies for overnight incubation. All primary antibodies and secondary antibodies used in this study are tabulated in Table S9 with RRID number. HRP-conjugated secondary antibody was incubated at room temperature for 1 h prior to signal development by Immobilon Forte Western HRP substrate (Sigma-Aldrich, #WBLUF0500). All images were captured with iBright CL1500 (Thermo Fisher Scientific, #A43678) and signal intensity was quantified with iBright Analysis software version 5.2.2.

### Fractionation assay

To determine the subcellular localization of BTNL9, we performed fractionation assay as described in a reference paper with minor modification(20). Briefly, 5 million cells were used for whole cell lysate (WCL), and isolation of cytoplasm fraction (Cyt) and nucleus fraction (Nuc). 15 µl of each fraction were loaded for western blot.

### Proliferation assay

Cell proliferation was determined by seeding 500 cells/well for A549 cell line and 150 cells/well for NCI-H460 with triplicates wells in 96-well microplate (Corning, #3596). One tenth of growth medium’s volume Resazurin (R&D Systems, #AR002) was added from Day 0 onward and incubated for 4 h at 37 °C in the presence of 5% CO_2_. Fluorescence was measured using microplate reader Viroskan Lux (Thermo Fisher Scientific, #VLBL00D0), by setting excitation at 544 nm and emission at 590 nm in Skanlt software 6.0.1 (Thermo Fisher Scientific). Multiple points per well were selected to eliminate cross-emission between wells. Growth curves were then generated by subtracting blank readings and normalizing to the initial reading on Day 0.

### Cytotoxicity assay

To assess the effect of BTNL9 overexpression on drug sensitivity, A549 and NCI-H460 cells were infected with lentivirus carrying either a vector control or BTNL9 overexpression construct. The following day, cells were counted and seeded at a density of 5,000 cells per well in 96-well plates in triplicate for each condition. On Day 2, either etoposide (Sigma-Aldrich, #E1383-25mg) or bortezomib (Sigma-Aldrich, #5043140001) were added to the wells at a range of concentrations to generate dose-response curves, and control wells were treated with vehicle only. Cells were incubated with the drugs for 48 hours at 37°C in a humidified incubator with 5% CO_2_. After the incubation period, resazurin solution was added and proceed followed the same as proliferation assay. The fluorescent signal was normalized to the vehicle control, and dose-response curves were plotted using PRISM software. The IC50 values were calculated for both vector control and BTNL9-overexpressing cells to evaluate the impact of BTNL9 on drug sensitivity.

### Cell cycle assay

To assess cell proliferation in a cell-cycle dependent manner, transfected cells were seeded in a 6-well microplate (NEST, #703003). After 24 h cells were pulsed with 10 µM of BrdU (Sigma-Aldrich, #B5002) for 1 h, harvested, and then stained using BrdU cell proliferation assay kit (BD Bioscience, #556028). Flow cytometry was performed using CytoFlex (Beckman Coulter, #A001-1-1102). The Flow cytometry data was analyzed by FlowJo software (version 10.2, RRID:SCR_008520). This experiments were repeated twice.

### Clonogenic assay

Initially, we determined the plating efficiency by seeding varying numbers of cells. For A549, 500 cells were seeded and for NCI-H460, 250 cells were seeded per 6 cm plate, experiments done in biological replicates and repeated twice (SPL Life Sciences, #20060). Plates were incubated at 37 °C in the presence of 5% CO_2_ for 10 days until colonies became visually apparent. The plates were stained with 0.1% crystal violet solution containing 20% of methanol for 20 min and gently rinsed with running water. Plate images were captured and colonies were counted with ImageJ (National Institutes of Health, RRID:SCR_003070).

### Soft agar assay

To determine anchorage-independent growth, soft agar assay was performed twice with biological triplicates. Low-gelling temperature agarose (Sigma-Aldrich, #A9414-25G) was dissolved in ultra-sterile water to make 2% agarose stock gel. The gel was liquified by heating and kept in a water bath set at at 45 °C. The plate was prewarmed inside incubator at 37 °C and coated with a 0.6% diluted gel to create a bottom layer, which was then solidified at 4 °C for 30 min. The base-coated plate was pre-warmed again at 37 °C and an adequate cell number (10,000 cells for A549, and 600 cells for NCI-H460 per well) was added into the 0.3% agarose containing growth media. The suspension was mixed gently avoiding bubbles formation and dispensed evenly on top of the base layer. The plate then was incubated at 4 °C for an additional 30 min, after which 1.5 ml of media was added to prevent the agar from disintegrating. After 2-3 weeks, colonies were stained by 1mg/ml of Nitro blue Tetrazolium Chloride (NBT) (Thermo Fisher Scientific, #N6495) at 37 °C overnight. Captured images were analyzed by OpenCFU (21).

### Cell Viability assay

To assess cell viability in BTNL9-overexpressing cells, a cell viability assay was employed to evaluate the impact of BTNL9 on cell death. A549 and NCI-H460 cells were seeded in a 6-well microplate (NEST, #703003) before infected with lentivirus carrying an empty vector control and pCMV-BTNL9. Subsequently, these cells were cultured for 3 days. Afterward, the cells were harvested, washed once with 1X MACS wash buffer (Miltenyi Biotec, #130-092-987), and then directly stained with eBioscience™ Fixable Viability Dye eFluor™ 780 (Thermo Fisher Scientific, #). We used 0.5 µl directly from the stock to stain each sample and incubated them on ice for 30 minutes while protecting them from light. After staining, the cells were washed twice with 1X MACS wash buffer and re-suspended in 100 µl of the same buffer. Flow cytometry analysis was performed using CytoFlex (Beckman Coulter, #A001-1-1102), and the flow cytometry data were analyzed using FlowJo software (version 10.2, FlowJo, RRID:SCR_008520). This experiments were repeated twice.

### Immunohistochemistry staining

IHC was conducted on paraffin-embedded tissue of normal lung tissue and lung tumor. We obtained unstained slides from Inje Biobank of Inje University Busan Paik Hospital, a member of Korea Biobank Network with Institutional Review Board regulations and approval (IRB #2023-09-012). Slides were heated for 1 h at 60 °C, deparaffinized in xylene, rehydrated in graded alcohol and rinsed in distilled water. Antigen retrieval and immunohistochemistry were performed using either a Benchmark XT or Discovery XT automated immunohistochemistry system (Ventana Medical Systems, Inc., Tucson, AZ) with an OptiView DAB IHC Detection kit. Slides were incubated for 34 min at room temperature with diluted BTNL9 antibody (LS-Bio; #LS-C399149). After antibody staining, sections were washed in distilled water, lightly counterstained with hematoxylin, rehydrated and mounted with coverslips. IHC images were obtained on a NanoZoomer RS Digital Pathology System scanner. The intensity of cytoplasmic staining was evaluated as low, intermediate and high grade.

### In vivo xenograft mouse model

4-weeks old male BALB/C nude mice were used for the subcutaneous tumor growth assay. Mice were housed in a 12-h light/12-h dark cycle at 22 °C and 50–60% humidity. Each mouse was subcutaneously injected in the flank with 100 µl of A549 transduced cells (5×10^6^) suspended in Matrigel Basement Membrane Matrix (BD Bioscience, #356234). Mice were randomly grouped into two (BTNL9 overexpression and empty vector control), each including 6 mice. The experiment was repeated twice (yielding a total of 12 mice per group) to minimize in vivo bias and outliers. Prior to the in vivo study, a plasmid expressing firefly luciferase marker was transfected into A549 cells, which were then selectively cultured with the antibiotic hygromycin B. Once a stable cell line was established, these cells were transduced with either empty vector control or BTNL9-OE lentivirus. At post-transduction 3 days, we implanted these cells subcutaneously into BALB/c nude mice. Tumor volume was measured in every 3 days by an investigator blinded for both experimental groups. Tumor volume was calculated as V = (LxW2)/2 where L is tumor length and W is tumor width, in millimeters (mm). The imaging was conducted and captured by In Vivo Imaging Systems (IVIS) at the start (Day 0) and end of the experiment (Day 30) using 30 mg/ml of D-Luciferin (GoldBio, #LUCK-1G). Mice were euthanized using CO_2,_ the tumors were excised for visual and weight-based assessments.

### Statistical analysis

All experiments were carried out independently three times. Data were expressed as mean ± standard deviation (SD) and compared using Student’s *t*-test and two-way ANOVA. Data with a *p*-value of *p* < 0.05 were considered a statistically significant difference (\**p* < 0.05, \*\**p* < 0.01, and \*\*\**p* < 0.001) was considered statistically significant. Data were processed with Graph PadPrism for Windows, version 8 (Graph Pad Software Inc., USA).

## RESULTS

### Characterization of protein structure and subcellular localization of BTNL9

Based on the annotated reference protein sequence deposited in the NCBI database (NP_689760.2), BTNL9 contains a basic leucine zipper (bZIP) domain in the region between 281 and 317. Notably, BTNL9 only contains a single leucine heptad repeat that deviates from the typical bZIP pattern of transcription factors, which usually display 4 to 5 heptad repeats. The positions “a” and “d” in the heptad repeat are important for dimerization stability and are typically occupied by hydrophobic amino acids which is not seen in BTNL9 protein sequence (Fig. 1A). Moreover, unlike other members of the bZIP transcription factor family, BTNL9 shows less similarity in the basic region with other transcription factors. These observations indicate that BTNL9 does not align with the typical characteristics of bZIP transcription factor profile, and it could potentially belong to a distinct type of transcription factor family.

**Fig. 1.**
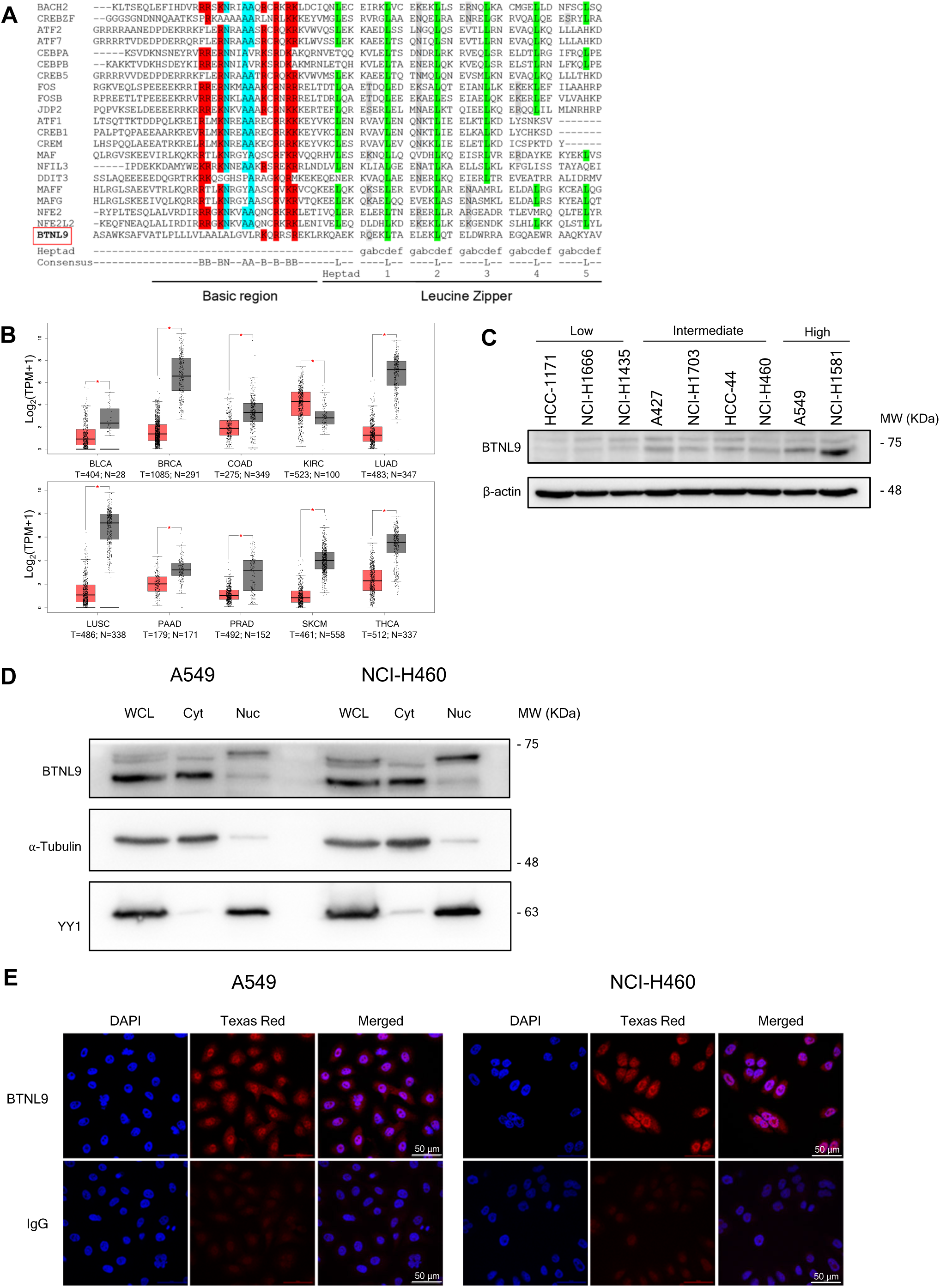
Characterization of protein structure and subcellular localization of BTNL9. **A.** Alignment analysis of bZIP transcription factor genes shows the conserved leucine zipper region, with BTLN9 corresponding to a single heptad repeat. Key annotations: conserved leucine (L) residues are highlighted in green, polar or charged amino acids in grey, basic amino acids in red, and conserved amino acids in the basic region in aqua blue. Consensus sequence is shown at the bottom of the figure, with B representing either arginine (R) or lysine (K). Heptad repeats and their positions are denoted as 1-5 and “abcdefg”, respectively. **B**, Boxplots depict the expression levels of BTNL9 (log_2_ TPM + 1) in tumor (T, red) and adjacent normal (N, gray) tissues across multiple cancer types, including BLCA (Bladder Urothelial Carcinoma), BRCA (Breast invasive carcinoma), COAD (Colon adenocarcinoma), KIRC (Kidney renal clear cell carcinoma), LUAD (Lung adenocarcinoma), LUSC (Lung squamous cell carcinoma), PAAD (Pancreatic adenocarcinoma), PRAD (Prostate adenocarcinoma), SKCM (Skin Cutaneous Melanoma) andTHCA (Thyroid carcinoma). The number of tumor (T) and normal (N) samples analyzed for each cancer type is indicated below each boxplot. Asterisks (*) indicate statistically significant differences in BTNL9 expression between tumor and normal tissues (*p < 0.05). This data were derived from TCGA dataset and figure was generated from GEPIA2. **C**, Western blot analysis of the protein levels of BTNL9 across different lung cancer cell lines. **D**, Subcellular fractionation of A549 and NCI-H460 cells followed by SDS-PAGE (8% gel) and western blot shows BTNL9 proteins in both cytoplasmic and nuclear fractions. Anti-⍺-tubulin and anti-YY1 were used to confirm the accurate partitioning of cytoplasmic and nuclear fractions, respectively. WCL: whole cell lysate, Cyt: cytoplasm, Nuc: nucleus. **E**, BTNL9’s localization within A549 and NCI-H460 were detected by immunofluorescence using rabbit anti-BTNL9 (Texas Red) or rabbit IgG isotype control, with DAPI (blue) for a nuclear counterstain. Confocal microscopic images were captured with 20X objective lens.

Before exploring the possibility of BTNL9 serving as a transcription factor, we conducted data mining using the TCGA dataset to assess BTNL9 mRNA levels across various cancer types and their normal counterparts. Overall, BTNL9 was found to be downregulated in tumor tissues compared to normal tissues (Fig. 1B). Given the prominent BTNL9 transcript expression in normal tissues and its notable disparity in tumor tissues, we selected the lung as the model for this study. Next, we examined a panel of human lung cancer cell lines for BTNL9 protein expression and chose the lung adenocarcinoma (LUAD) cell lines A549 and NCI-H460 due to their relatively rapid doubling time and median BTNL9 expression levels for all the subsequent in vitro experiments (Fig. 1C). We did not use the NCI-H1581 cell line because it is a semi-suspension cell line that is not suitable for certain phenotypic studies.

Subsequently, we evaluated the subcellular localization of BTNL9 through protein fractionation followed by western blot analysis to delineate its expression profile (Fig. 1D). The whole cell lysate analysis of A549 and NCI-H460 revealed three bands for BTNL9, each below 75 KDa, with an estimated molecular weight (MW) of 60 KDa based on the full-length BTNL9 amino acid (aa) composition. Notably, the largest protein mainly localized in the nucleus, while the other smaller ones were mostly detected in the cytoplasm. Utilizing NLStradamus program for nuclear localization sequence (NLS) prediction (22), we found that BTNL9 possesses an NLS at positions 278-294 aa (Fig. S1A). Protein sequence alignments of BTNL9’s various transcript variants showed a conserved NLS domain, except for the variants X8 and X9, which have markedly lower MWs (Fig. S1B). Based on their higher MWs, the three BTNL9 bands shown in the western blot in Fig. 1D are likely to have an NLS for nuclear translocation. Yet, it remains unclear why only the BTNL9 isoform with the highest MW was located in the nucleus, with the remaining two isoforms staying in the cytoplasm.

To visually validate the nuclear presence of BTNL9, we conducted confocal microscopy on A549 and NCI-H460 cells. Consistent with the protein fraction results, BTNL9 was observed in both the cytoplasmic and nuclear compartments, with a pronounced nuclear localization in both cell lines (Fig. 1E). This pattern of distribution aligns with the typical localization characteristics of transcription factors.

### Insight of BTNL9 genome-wide binding sites

In order to investigate the DNA-binding capacity of BTNL9, we performed chromatin immunoprecipitation sequencing (ChIP-seq) in A549 cells. Due to the low BTNL9 protein level in A549 cells, we overexpressed BTNL9-AM-tag plasmid and ChIP were pulldown with AM-tag antibody. The ChIP-seq data revealed a wide distribution of genome binding sites: while 10% of the sites were detected in the upstream promoter and exon regions, the majority (∼87%) were located within intergenic and intronic regions (Fig. 2A). The similar distribution of DNA-binding profiles have been found in other transcription factors (23, 24). The density plot of read counts and heatmap analysis indicated that BTNL9 bound to transcription start site (TSS) regions, where these genes were actively transcribed (with strong peaks observed for H3K27ac) (Fig. 2B and C). This provides initial evidence of the ability of BTNL9 to bind to promoter regions. Utilizing MEME-ChIP motif analysis, we deciphered the potential conserved DNA binding motif of BTNL9 (Fig. 2D and Fig. S2A). The conserved sequence CCCCRCCCC was derived from the central 200 bp position of BTNL9 ChIP-seq peak replicates data.

**Fig. 2.**
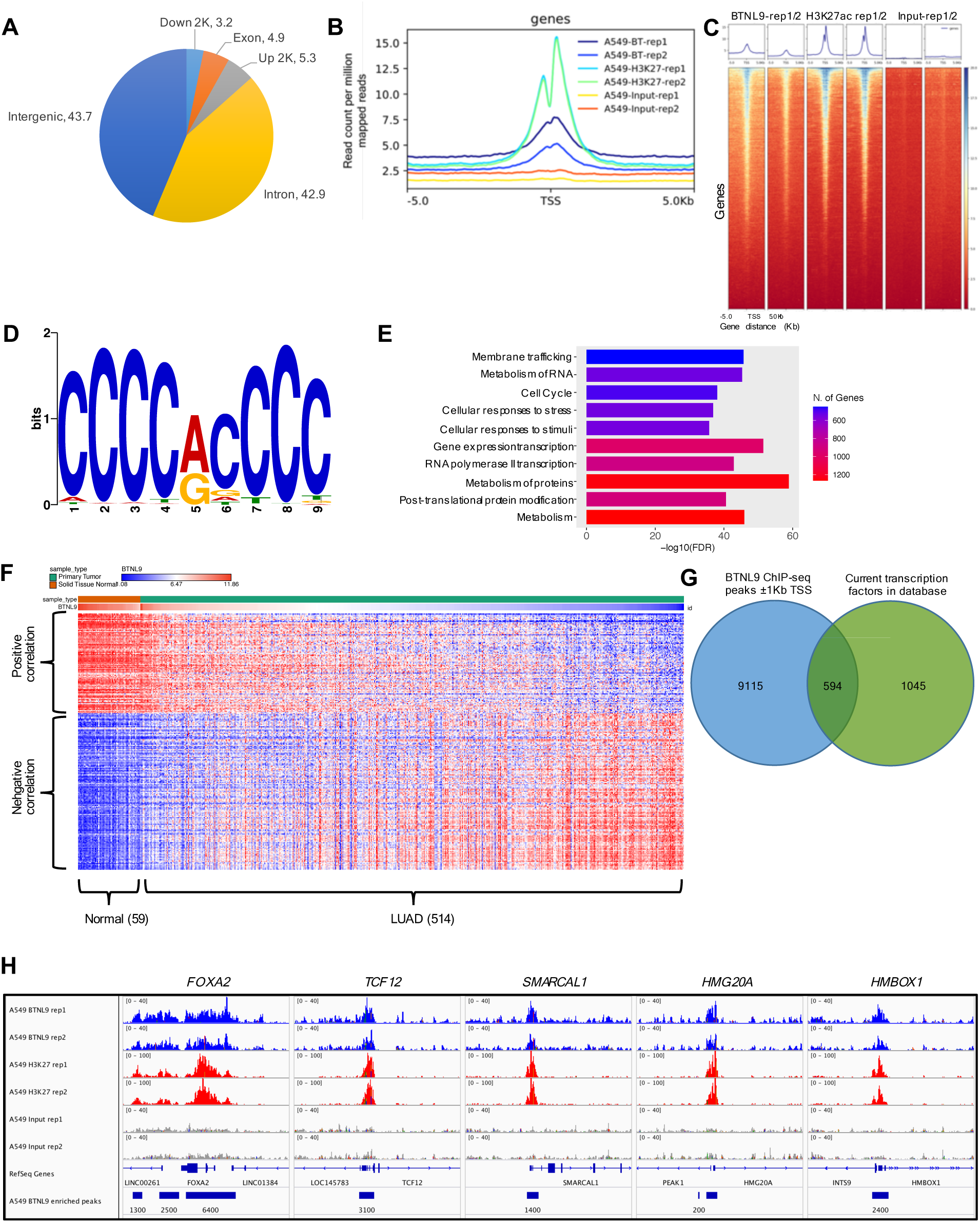
Insight of BTNL9 genome-wide binding sites. **A**. Distribution of BT NL9 binding sites based on pe ak location across different genomic regions of the human genome. **B**, Density plot of read counts showing the relative ChIP-seq read de nsity near the transcriptional start site (TSS) of BTNL9, H3K27ac and ChIP input within ± 5 Kb. **C**, H eat map illustrating the quantity of genes bound by BTNL9, H3K27ac and ChIP input control arou nd TSS. **D**, *De novo* motif of the potential conserved DNA binding motif of BT NL9 was generated by us ing MEME-ChIP motif analysis. **E**, Bar graph representing Reactome pat hway analysis of gen es identified from ChIP-seq differential peak calling. The top 10 significant pathways were shown. **F**, Heatmap representing expression profiles of ge nes significantly correlated to BTNL9 mRN A levels from TCG A LUAD dat aset (Spearman’s correlation coefficient *r* ≤ – 0.45 or *r* ≥ 0.45). **G**, The Venn diagram displays the overlap betw een gen es identified by C hIP-seq and the complete list of transcription factors, highlighting potential transcription factors directly regulated by BTNL9. **H**, Integrative Genomic View er (IGV) pot displays ChIP-seq peak of BTNL9, H3K27ac, and input control at the loci of the top 5 transcription factors in the list.

Through ChIP-seq peak calling, we filtered peaks based on *p* < 1.0E-5, following MACS recommendations, with a false discovery rate (FDR) of *q* < 0.005. A total of 26,610 peaks were selected for further analysis (Table S1). We then used the Genomic Regions Enrichment of Annotations Tool (GREAT) to annotate ChIP peaks to the nearest genes within ±1 kb of the transcription start site (TSS), identifying 9,707 genes associated with BTNL9 binding sites (Table S2). To validate these findings, we randomly selected three genes (*DDX1*, *UBC*, and *KANSL3*) (Fig. S2B) and performed ChIP-qPCR in both A549 and NCI-H460 cells overexpressing BTNL9-AM-tag (Fig. S2C). Using the percent of input method, we observed significant enrichment of ChIP-DNA at these gene loci across both cell lines (Fig. S2C).

We analyzed these 9,707 genes using the ShinyGo V0.82 tool to assess Reactome pathway enrichment, identifying the top 10 enriched functional pathways: membrane trafficking, metabolism of RNA, cell cycle, cellular responses to stress and stimuli, gene expression transcription, RNA polymerase II transcription, metabolism of protein, post-translational protein modification and metabolism. (Fig. 2E). The results of this analysis imply that BTNL9 binds to genes critical for gene regulation and cell stress responses. To investigate the potential transcriptional regulation of BTNL9 in lung cancer, we performed Spearman’s correlation analysis between BTNL9 and these 9,707 genes RNA-seq data in lung adenocarcinoma (LUAD) and lung squamous cell carcinoma (LUSC) patients. Using cBioPortal’s TCGA database, we identified 249 genes with substantial clinical relevance to BTNL9; among them, 92 genes were positively correlated and 157 were negatively correlated, based on Spearman’s correlation coefficient cutoffs of *r* ≤ –0.45 or *r* ≥ 0.45 (Table S3). Heatmaps were generated to visualize the correlation patterns in LUAD (59 normal, 514 tumor samples) and LUSC (51 normal, 502 tumor samples) (Fig. 2F and Fig. S2D).

In normal tissues adjacent to LUAD, genes formed distinct clusters of positive and negative correlations, suggesting that BTNL9 may have a differential regulatory role. In LUAD tumor samples, the correlation patterns followed a gradient based on BTNL9 expression levels, indicating that these genes may be regulated in a BTNL9-dependent manner. In contrast, LUSC samples exhibited a more asymmetric correlation pattern. While positively correlated genes aligned with BTNL9 expression levels, negatively correlated genes in LUSC did not show a clear relationship with BTNL9, suggesting a weaker or more indirect regulatory influence. This contrast between LUAD and LUSC implies that BTNL9 may have distinct transcriptional roles in different lung cancer subtypes.

We next examined whether BTNL9 specifically binds to the promoter or intronic regions of any potential transcription factors. To this end, we compared 9,707 genes from our ChIP-seq data with an updated list of 1,639 transcription factors from the AnimalTFDB V4 database. Through this analysis, we identified 594 transcription factors that BTNL9 binds to (Fig. 2G and Table S4). We highlighted five transcription factors (*FOXA2*, *TCF12*, *SMARCAL1*, *HMG20A* and *HMBOX1*), where BTNL9 binding sites significantly overlapped with H3K27ac peaks, indicative of active transcription regions (Fig. 2H). These findings strongly imply that BTNL9 binds to a substantial number of genes either at the promoter or in intronic regions, highlighting its potential role as a novel transcription factor in a multilayered hierarchical gene regulatory networks.

### Analysis of BTNL9 RNA-seq data obtained in A549 cell line

We investigated the role of BTNL9 in A549 cells by using lentivirus to induce overexpression of BTNL9 with an empty vector or a BTNL9-overexpression (BTNL9-OE) vector. On Day 3, total RNA was harvested for RNA-seq. Our preliminary data showed that BTNL9 expression in A549 reached its peak levels on Day 3, after which it gradually decreased (Fig. S3A). We sent duplicate total RNA samples from each experimental group to E-biogen (South Korea) for RNA-seq. For further analysis, we selected 444 differentially expressed genes (DEGs) using cutoff values of Log_2_ FC ± 0.5 and FDR *q* < 0.1. The MA plot revealed an even distribution of DEGs, with 176 genes identified as upregulated and 268 genes as downregulated (Fig. 3A and B, Table S5). To elucidate the functions of BTNL9, we employed the ShinyGo V0.82 tool for DEG analysis. This analysis included Reactome pathway enrichment, which highlighted the top 10 principal pathways related to the regulation of the cell cycle and cell division (Fig. 3C). Additionally, we generated a heatmap with hierarchical clustering of genes influenced by BTNL9 expression and detected three clusters that were associated with DNA replication, transcription regulation mediated by p53, and cell cycle (Fig. 3D).

**Fig. 3.**
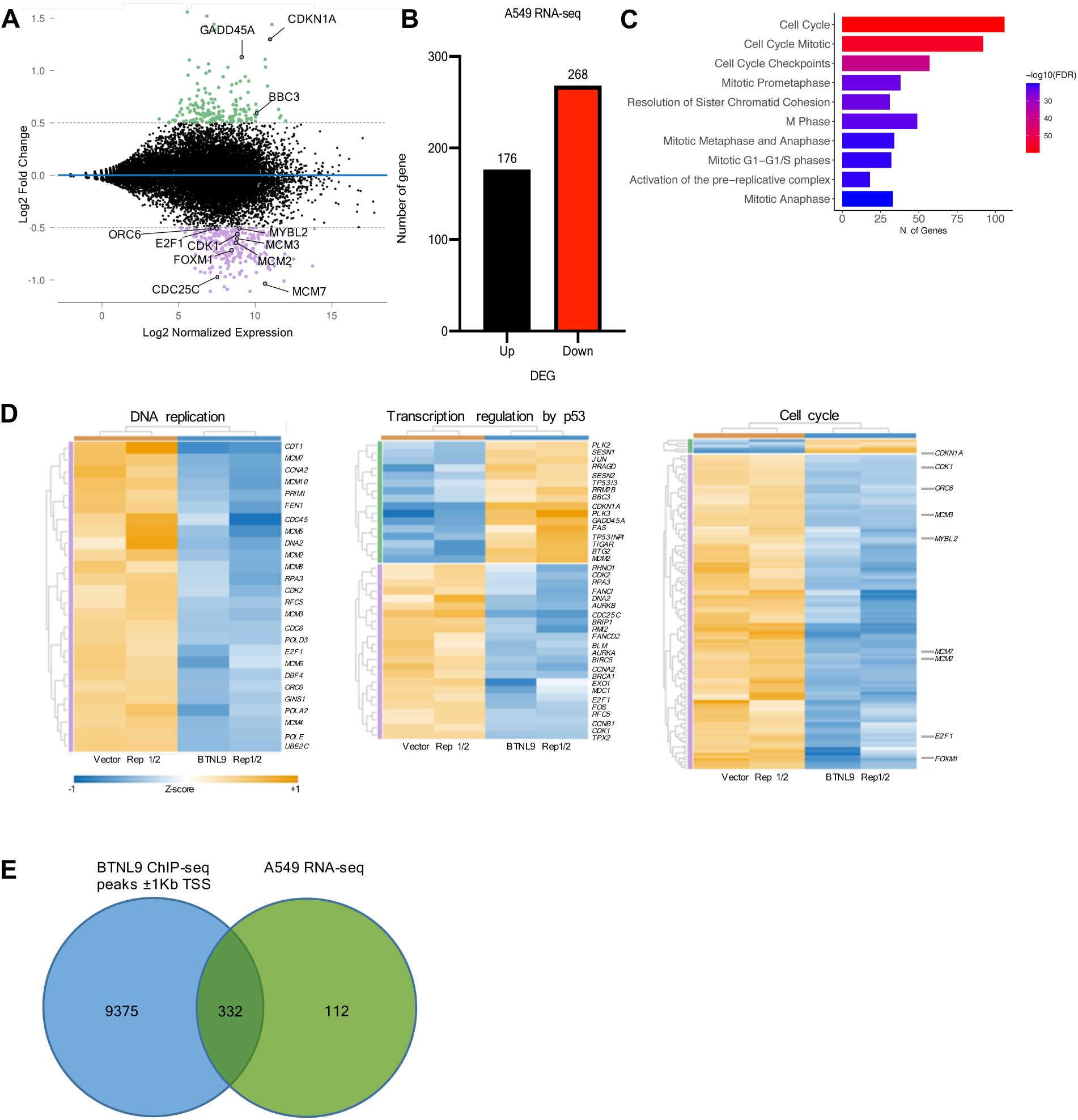
**Regulatory role of BTNL 9 in cell cycle, DNA replication, and p 53-regulated genes in lung cancer**. **A**. MA plot displaying the distribution of D EGs from A549 RNA-seq dataset, with upregulated gen es shown in green and downregulated genes in purple. This analysis is based on the cut-off value of Log_2_ FC ± 0.5 and an false discovery rate (FDR) of < 0. 1. Genes marked with labels were further analyzed in subsequent experiments. **B**, Bar graph showing the number of D EGs that met the criteria of Log_2_ FC ± 0.5 and FDR < 0.1. **C**, Bar graph representing the top 10 significant pat hways from the Reactome pathway analysis of statistically significant genes identified from the RNA-seq. **D**, Heatmap generated using the PAM (Partitioning Around Medoids) method with the fpc R library, showcasing 3 major DEG clusters associated with DNA replication (26 gen es), transcription regulation by p53 (36 genes) and cell cycle (119 gen es). **E**, Venn diagram showing the overlap between two sets of genes identified through both ChIP-seq and RNA-seq analyses in A549 cells.

Within the DNA replication cluster regulated by BTNL9, it was evident that the entire set of minichromosome maintenance protein (MCM) family genes, including *MCM2* through *MCM8* and *MCM10*, were downregulated (Table S6). The MCM complex acts as a DNA helicase and is a crucial component of the DNA replication licensing system (25). Moreover, we found that BTNL9 regulates a range of genes critical for DNA replication. These include licensing factors (*CDT1*), genes involved in the G2/M transition (*CCNA2*, *CDK2* and *UBE2C*), and polymerase catalytic subunits (*POLE* and *POLA2*), as well as DNA primase that synthesizes small RNA primers for Okazaki fragment synthesis (*PRIM1*).

We also identified a cluster of 36 genes among the DEGs affect by BTNL9 that are involved in p53-mediated transcription regulation (Table S6). These genes modulate various aspects of cellular activity such as cell cycle checkpoint, DNA repair, DNA replication, and programmed cell death. Critical effectors in this cluster include *BBC3*, *CDC25C*, *CDK1*, *CDKN1A*, *E2F1*, *GADD45A* and *MDM2*. This finding is particularly interesting because it implies that BTNL9 co-regulates a subset of genes related to p53-mediated modulation, thereby raising the possibility that it plays a role in the safeguarding of genome integrity.

Our analysis of the cell cycle cluster demonstrated that a large number of cell cycle-related genes (119 genes) were regulated by BTNL9, and the majority of these genes were downregulated (Table S6). These genes include common cell cycle regulators such as cyclins and CDKs, cell division cycle genes, and genes encoding centromere-associated proteins such as *CENPA* and *CENPE* (Table S6). Inactivation of these genes is known to lead to mitotic arrest; moreover, CENPA plays an important role in kinetochore recruitment for chromatid segregation (26–28). Furthermore, genes encoding condensins, which are structural maintenance of chromosome (SMC) proteins, and non-SMC condensins were also identified within the cluster, crucial for maintaining sister chromatid structure during mitosis and meiosis (29).

We compared the 444 DEGs with the 9,707 genes identified from BTNL9 ChIP-seq analysis and found 332 overlapping genes (Fig. 3E). This indicates that 74.8% of the DEGs are directly regulated by BTNL9 (Table S7).

### Regulatory role of BTNL9 in cell cycle progression through multiple pathways

To gain a deeper understanding of BTNL9’s functions in cancer cells, especially in DNA replication, cell cycle and p53 mediated regulation, we selected several well-characterized genes that could elucidate its role in suppressing tumor activities. We focused on 12 genes from the three clusters identified in our RNA-seq data, as their potential to inhibit tumor growth was suggested by their overlapping functions (Fig. 4A). String analysis confirmed that these genes were closely linked to each other (Fig. S4A). Furthermore, our ChIP-seq data provided direct evidence of BTNL9 binding to some of these genes, either at the TSS or within intronic regions. From our ChIP-seq analysis, we determined that 8 genes, *BBC3, CDC25C, CDK1, FOXM1, GADD45A, MCM3, MCM7 and ORC6* exhibited significant ChIP-seq peaks (Fig. S4B and Table S2). This suggests that BTNL9 acts as an upstream transcription factor, either as an enhancer or repressor, regulating the expression of these genes. Subsequently, we validated the RNA expression profiles of all 12 genes using RT-qPCR in both A549 and NCI-H460 cell lines, specifically from Day 2 to Day 4 following the infection with BTNL9 lentivirus. Consistent with the RNA-seq data, we noted significant changes in gene expression on Day 3, which became more pronounced on Day 4 in both cell lines (Fig. 4B and Fig. S4C). Among these genes, *BBC3*, *CDKN1A* and *GADD45A* were significantly upregulated, whereas *CDC25C*, *CDK1*, *E2F1*, *FOXM1*, *MCM2*, *MCM3*, *MCM7*, *MYBL2,* and *ORC6* showed consistent downregulation in both cell lines.

**Fig. 4.**
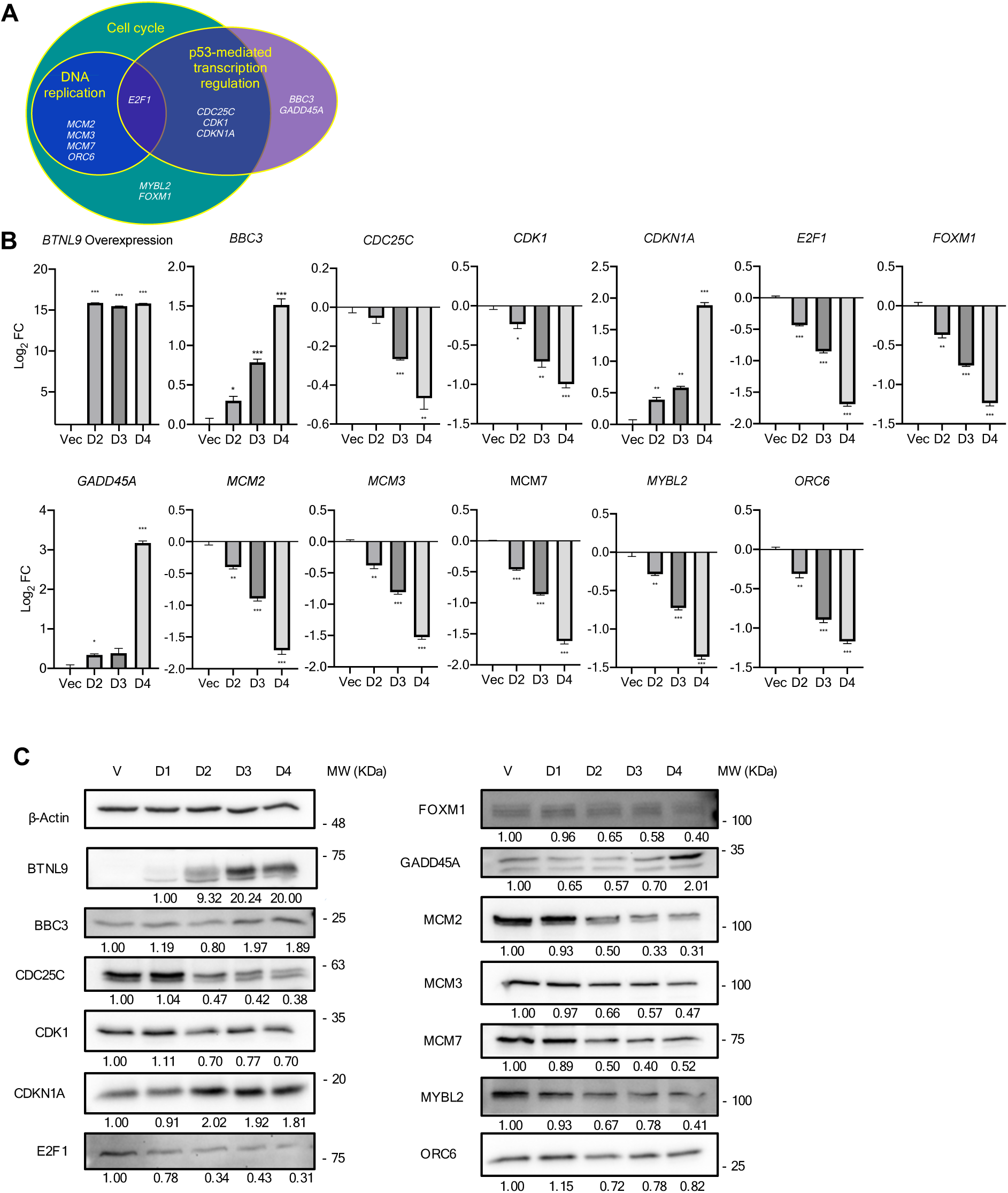
Validation of RNA-seq data using qRT-PCR and western blot A, Venn diagram illustrating the over lap between the functional pathways of the selected genes.. **B**, Bar graphs representing qRT-PC R analyses in A549 cells overexpressing either empty vector control or BTNL9, with samples harvested on the indicated day. All values were calculated as mean Log_2_ FC ±SD. Significance was determined by a two-tailed Student’s *t*-test: * *p* < 0.05, ** *p* < 0.01 and *** *p* < 0. 001. All experiments were conducted in triplicates. Primers for qRT-PC R of tested genes w ere listed in Table S8. **C**, Representative western blot of selected gen es in A549 cells overexpressing either empty vector control or BTNL9, with samples harvested on the indicated day. β-actin was used as a loading control. Signal intensities w ere quantified with iBright Analysis software and semi-quantitative protein expression levels were calculated by normalizing to β-actin.

To determine the protein levels of these genes, we harvested protein lysates from cells infected with either BTNL9-or vector-lentivirus at specified time points and performed western blot for the same genes that were previously examined using RT-qPCR. In both cell lines, the trends in protein expression of most genes were consistent with those observed at the mRNA level (Fig. 4C and Fig. S4D). This result suggests that the expression levels of BTNL9 were sufficient to drive the transcription and translation of its downstream targets. Subsequently, we aimed to clarify whether p53 was involved in the regulation of BTNL9-regulated DEGs, particularly those within the p53-regulated gene cluster. To elucidate the relationship between BTNL9 and p53, we generated p53 knockout (KO) cell lines with CRISPR-cas9 technique in both A549 and NCI-H460 cells. Western blot analysis confirmed the successful knockout, as no p53 protein expression was detected in p53 KO cells, even in the presence of Nutlin-3, a known MDM2 inhibitor (Fig. S4E).

To investigate the p53-dependent and p53-independent regulatory effects of BTNL9, RT-qPCR was performed to analyze the expression of several BTNL9 downstream target genes (*BBC3*, *CDC25C*, *CDK1*, *CDKN1A*, and *E2F1*) previously shown to be associated with p53 regulation (Fig. S4F). In wild-type (WT) cells overexpressing BTNL9, *BBC3* and *CDKN1A* expression were markedly upregulated, while *CDC25C*, *CDK1*, and *E2F1* were downregulated, consistent with p53-mediated regulation. However, in p53 KO cells, BTNL9 overexpression failed to induce significant changes in these 5 genes expression. These findings highlight the role of BTNL9 in modulating downstream targets in a p53-dependent manner.

### Suppressive effect of BTNL9 on cell proliferation, clonogenicity, and tumorigenicity in lung cancer cells

Having demonstrated that BTNL9 regulates the expression of a group of genes involved in cell cycle progression, we aimed to investigate cellular phenotypes, such as cell proliferation and clonogenicity, to further support BTNL9’s regulatory role in cancer cells. Through the cell proliferation assay using A549 and NCI-H460 cell lines, we observed a significant inhibition of cell proliferation on Day 5 in BTNL9-OE cells compared to the empty vector-expressing control cells in both cell lines (Fig. 5A). To gain further insights into the mechanisms underlying this decline in proliferative activity, we conducted BrdU labeling assays to investigate whether this effect was cell cycle-dependent or associated with cell death Our cell cycle analysis revealed dramatic cell cycle arrest at the G0/G1 phase, along with reduced replication activity during the S phase (Fig. 5B). Additionally, our clonogenic assay showed a decrease in colony formation in BTNL9-OE cells compared to the empty vector-expressing control cells (Fig. 5C). These findings were further supported by the results of cell viability assessments using eBioscience™ Fixable Viability Dye eFluor™ 780 staining analysis, which demonstrated a significant reduction in cell viability in cells overexpressing BTNL9 (Fig. 5D). To further investigate the impact of BTNL9 on tumorigenicity, we conducted an anchorage-independent assay, which revealed a drastic reduction in colony formation in BTNL9-OE cells (Fig. 5E). Overall, our experimental results provide compelling evidence for the regulatory role of BTNL9 in modulating both proliferative activity and tumorigenicity in LUAD cell lines.

**Fig. 5.**
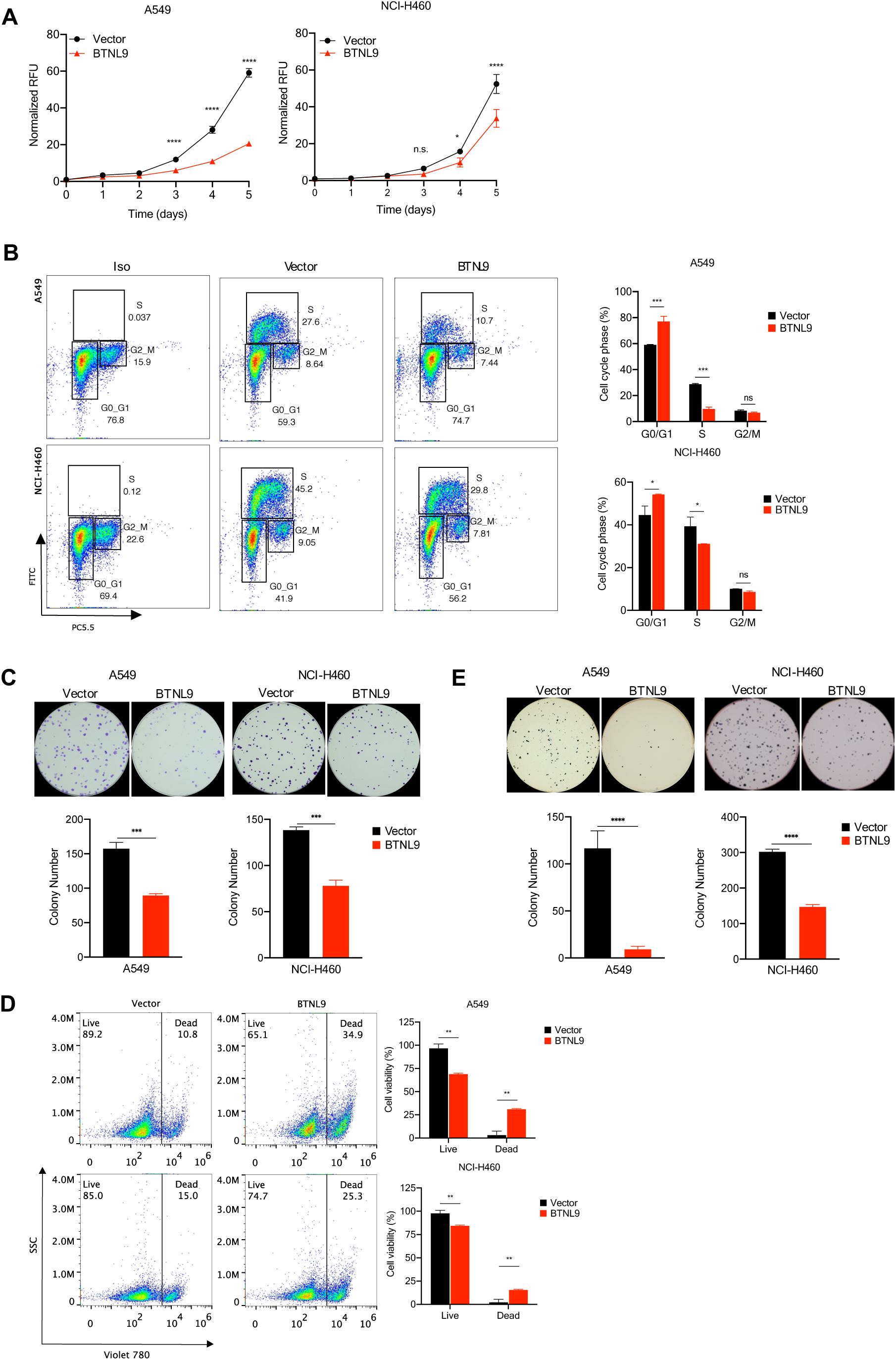
**Suppressive effect of BTNL9 on cell proliferation and tumorigenicity in lung ncer cells**. **A**, 49 and NCI-H460 cells, overexpressing either empty vector control or BTNL9, were used in the following assays. All experiments were conducted in biological replicates. **A**, Cell proliferation of both cells was evaluated using resazurin to measure cell growth over 5 consecutive days. Growth curves were plotted as mean relative fluorescence units (RFU) ± SD. **B**, Representative scatter plots for the cell cycle assay, performed using BrdU staining on both cells, with isotype control serving as a negative control. Cell populations representing G0/G 1 phase, S phase and G2/M phase w ere gated. Bar graphs representing the compilation of individual experimental results with mean ± SD. **C**, Clonogenic assay was conducted to investigate survival differences between an empty vector control and BTNL9 overexpression groups. The number of colonies was counted and the bar graph was plotted as the mean ± SD. **D**, Representative scatter plots for the cell viability assay, measured and stained using eBioscience™ Fixable Viability Dye eFluor ™ 780. Bar graphs displaying the live and dead populations, plotted as mean ± SD. **E**, Representative images of colonies from anchorage-independent growth assay between the empty vector control and BTNL9 overexpression groups. The number of colonies w as counted and plotted in bar graphs as mean ± SD. For **A**, **B** and **D** data represents the mean of triplicates and tested with two-w ay ANOVA using the Tukey test. \**p* < 0. 05, \*\**p* < 0.01, \*\*\**p* < 0.001, \*\*\*\**p* <0.0001, and ns denoting no statistical differences. For **C** and **E** data represents the mean of triplicates and tested by Student’s *t* test (unpaired two-tailed). \**p* < 0. 05, \*\**p* < 0.01, and *p* ***< 0.001

### In vivo demonstration of the tumor suppression effect of BTNL9

To investigate BTNL9’s tumor-suppressive effect under in vivo conditions, we used a xenograft mouse model by subcutaneously implanting A549 cells transduced with either BTNL9-or vector-lentivirus, and employed bioluminescence imaging to quantify tumor size. Images captured by IVIS on Day 30 revealed significantly larger tumors in the group implanted with A549 cells expressing the empty vector control cells than in the group implanted with BTNL9-OE cells (Fig. 6A). Tumor volumes were also measured every 3 days until Day 30, and notable differences between the groups were discernable from Day 18 onward (Fig. 6B). Both visual and weight-based assessments confirmed marked tumor growth inhibition in the group implanted with cells overexpressing BTNL9 (Fig. 6C). These findings indicate that BTNL9 act as a tumor suppressor protein.

**Fig. 6.**
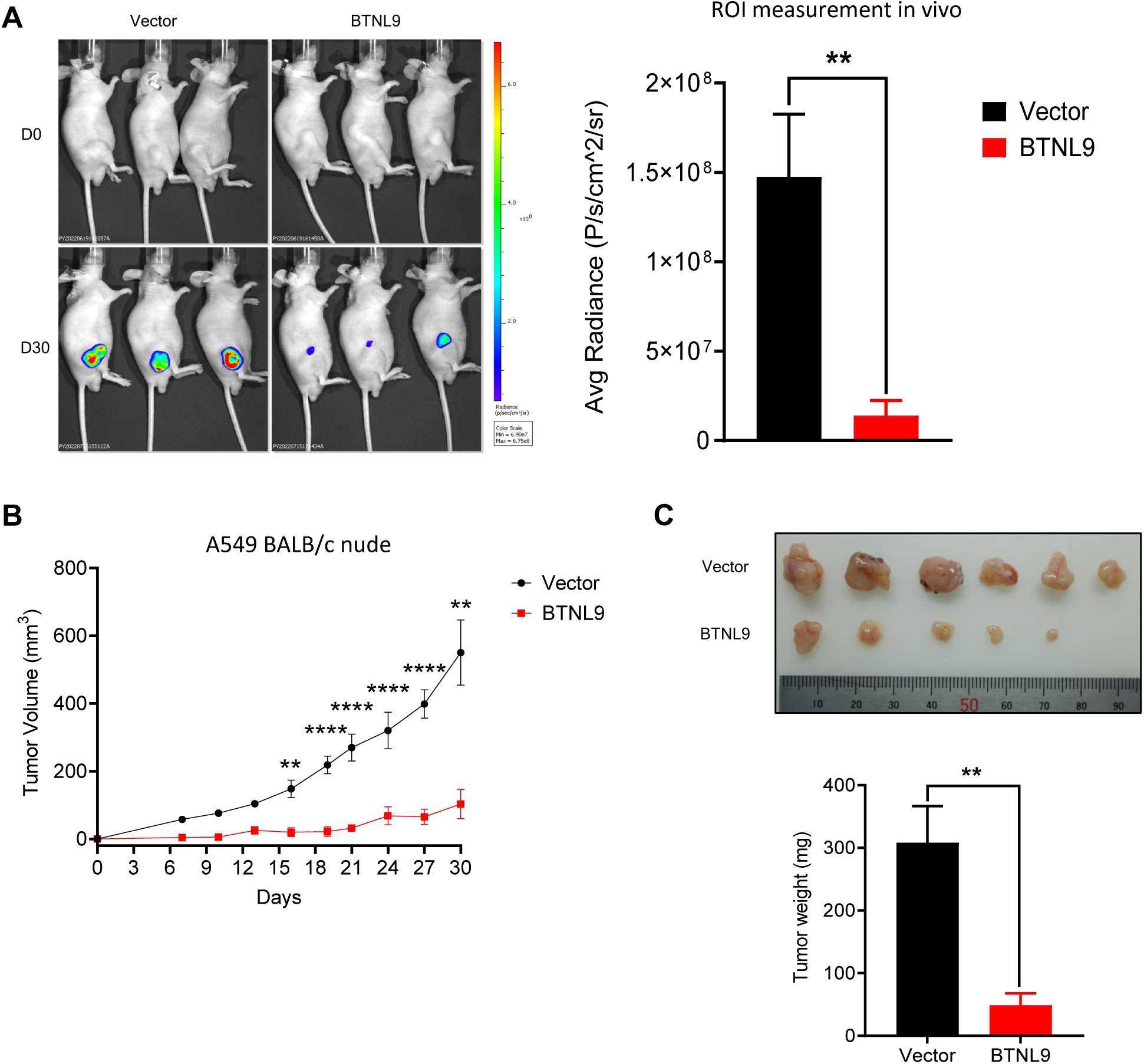
**In vivo demonstration of the tumor suppression effect of BTNL9**. **A**. BALB/c nude mice bearing subcutaneous A549 tumors in each group were imaged by IVIS on day 0 and day 30. The average radiance at the tumor site was normalized to the day 0 signal, and data were displayed in bar graphs as mean ± SEM (n = 6; only 3 mice in each group are shown). P values w ere calculated using a two-tailed student’s t-test, \*\**p* < 0.01. **B**, Tumor volumes in each group were measured every 3 days, tumor growth curve was plotted as mean ± SEM (n=6). A significant difference in growth rate was determined using two-tailed student’s t-test (\*\**p* < 0.01 and \*\*\*\**p* < 0.00 01). **C**, Ex vivo image of the solid tumors of each group was photograp hed and the weights were plotted in bar graph as mean ± SEM (n=6; \*\**p* < 0. 01; unp aired two tail t-test). In vivo experiments were repeated twice.

### Higher BTNL9 expression level enhances drug sensitivity in lung cancer cell lines

To investigate the role of BTNL9 in modulating drug sensitivity in lung cancer cells, we study the effects of BTNL9 expression level in response to commonly used chemotherapy drugs. Using cytotoxicity assays, cell cycle analyses, and downstream marker expression profiling, we aimed to elucidate how BTNL9 influences the efficacy of etoposide and bortezomib treatments.

To evaluate the role of BTNL9 in drug sensitivity, cytotoxicity assays were performed on A549 and NCI-H460 cells with either vector control or BTNL9 overexpression. BTNL9 overexpression significantly enhanced drug sensitivity in A549 and NCI-H460 lung cancer cell lines. In cells overexpressing BTNL9, the IC50 values for etoposide and bortezomib were reduced compared to vector control cells. Specifically, for etoposide, the IC50 decreased from 69.13 μM in vector cells to 57.74 μM in BTNL9-overexpressing A549 cells. Similarly, for bortezomib, the IC50 dropped markedly from 0.20 μM in vector cells to 0.01 μM in BTNL9-overexpressing A549 cells (Fig. 7A). A similar trend was observed in NCI-H460 cells. When treated with etoposide, the IC50 showed a slight reduction from 0.14 μM in vector cells to 0.06 μM in BTNL9-overexpressing cells. In contrast, bortezomib treatment led to a substantial decrease in IC50, dropping from from 9.30 μM in vector cells to 0.27 μM in BTNL9-overexpressing cells (Fig. S5A).

**Fig. 7.**
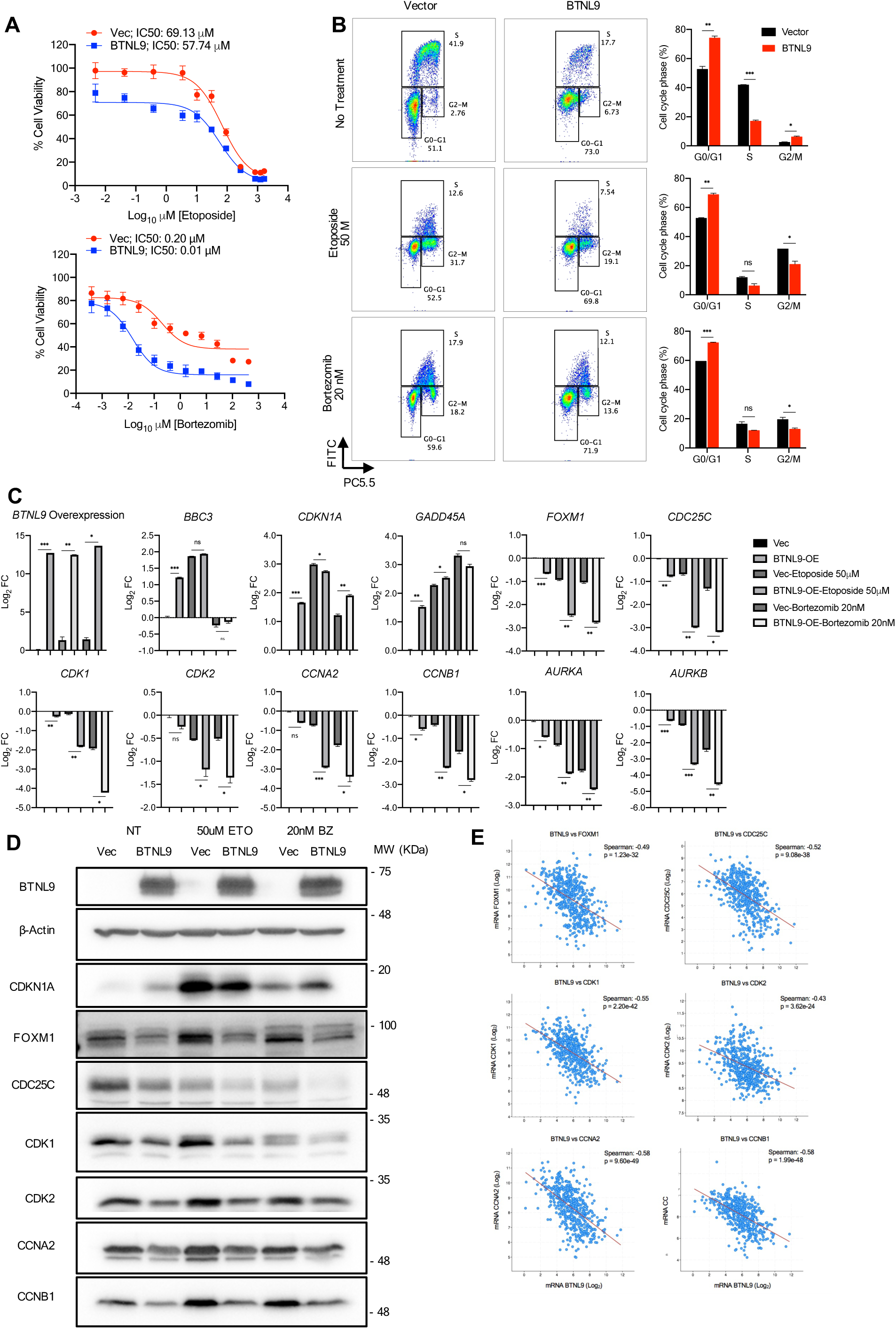
**BTNL9 enhances drug sensitivity to etoposide and bortezomib in lung cancer cells by regulating cell cycle pathways in A549**. **A**. Dose-response curves from cytotoxicity assays of et oposide and bortezomib treatments for 48 hr in A549 cells overexpressing BTNL9 or vector control was used to estimate for IC50 value. **B**, Representative scatter plots for the cell cycle assay, performed using BrdU staining on vector control and BTNL9-overexpressing in A549 cells. Cell populations representing G0/G1 phase, S phase and G2/M phase were gated. Bar graphs representing the compilation of individual experiment al results with mean ± SD. **C**, Bar graphs representing RT-qPCR analyses in A549 cells overexpressing either empty vector control or BTNL9. All values were calculated as mean Log_2_ FC ±SD. Significance was det ermined by a two-tailed Student’s *t*-test: * *p* < 0.05, ** *p* < 0.01 and *** *p* < 0.001. All experiments were conducted in biological triplicates. Primers for qRT-PC R of tested genes w ere listed in Table S8. **D**, Representative western blot of selected key cell cycle markers in A549 cells overexpressing either empty vector control or BTNL9. β-actin was used as a loading control. **E**, Correlation analysis betw een BTNL9 mR NA levels and the expression of cell cycle-related ge nes (*FOXM1*, *CDC25C*, *CDK1*, CDK2, *CCNA2*, and *CCNB1*) in LU AD patient samples derived from TCG A dat aset. Spearman correlation coefficients and p-values are shown, indicating a ne gative correlation between BTNL9 expression and these cell cycle-promoting genes. For **B**, **C**, and **D**, cells were either w ith no treatment (NT), or treated with 50 µM etop oside or 20 nM bortezomib for 18 hr. For **B** and **C** data represents the mean of biological triplicates and tested by Student’s *t* test (unpaired tw o-tailed). * *p* < 0.05, ** *p* < 0.01 and *** *p* < 0.001.

BrdU cell cycle assay showed that BTNL9 overexpression significantly affected cell cycle distribution, particularly under drug treatment conditions. Without treatment, BTNL9-overexpressing cells exhibited a higher proportion in the G0/G1 phase and a significantly lower percentage in the S phase compared to vector control cells. Upon treatment of BTNL9-overexpressing cells with 50 μM etoposide or 20 nM bortezomib for 18 hr, the percentage of NCI-H460 cells in the G0/G1 phase further increased, whereas little change was observed in A549. A concomitant significant reduction in S-phase cells compared to vector controls, along with further accumulation in G2/M phase were observed in both cell lines (Fig. 7B & S5B). This suggests that higher BTNL9 levels in cancer cells enhance G0/G1 phase arrest in NCI-H460 and impede cells progression in G2/M phase in both cell lines, potentially contributing to its efficacy in drug treatment.

RT-qPCR analysis of both A549 and NCI-H460 cells revealed that BTNL9 overexpression led to the downregulation of key cell cycle-related genes, including *FOXM1*, *CDC25C*, *CDK1*, *CDK2*, *CCNA2*, *CCNB1*, *AURKA* and *AURKB* particularly enhanced its suppression under drug treatment conditions. Whereas, *BBC3*, *CDKN1A* and *GADD45A* did not produce strong modulations compared to those key cell cycle related genes (Fig. 7C and Fig. S5C). Western blot analysis confirmed these findings at the protein level. BTNL9-overexpressing cells in A549 and NCI-H460 treated with 50 μM etoposide or 20 nM bortezomib for 18 hr exhibited reduced protein levels of FOXM1, CDC25C, CDK1, CDK2, CCNA2, and CCNB1. In contrast, CDKN1A expression was upregulated following treatment with both drugs, with a more striking difference observed between the vector control and BTNL9 overexpression groups, particularly in response to bortezomib (Fig. 7D and Fig. S5D).

Using RNA-seq data from TCGA dataset, correlation between BTNL9 expression and cell cycle-related genes in LUAD and LUSC were analysed. From LUAD patient dataset, Spearman’s correlation analysis revealed a significant inverse relationship between BTNL9 mRNA expression and key cell cycle-related genes. BTNL9 showed strong negative correlations with FOXM1 (Spearman: –0.49, *p* = 1.23e-32), CDC25C (–0.52, *p* = 9.08e-38), CDK1 (–0.55, *p* = 2.20e-42), CDK2 (–0.43, *p* = 3.62e-24), CCNA2 (–0.58, *p* = 9.60e-49), and CCNB1 (–0.58, *p* = 1.99e-48) (Fig. 7E). These correlations suggest that high BTNL9 expression is associated with the suppression of genes involved in cell cycle progression, particularly those required for S-phase and G2/M transition. From LUSC patient dataset, the negative correlations were weaker but remained statistically significant for most genes (Fig. S5E). For FOXM1, the correlation was weaker (Spearman: –0.10, *p* = 0.0232), and for CDC25C, no significant correlation was observed (Spearman: –0.07, *p* = 0.124). However, negative correlations were significant for CDK1 (–0.21, *p* = 2.01e-6), CDK2 (–0.10, *p* = 0.0266), CCNA2 (–0.24, *p* = 6.18e-8), and CCNB1 (–0.23, *p* = 1.56e-7). This suggests that the relationship between BTNL9 and cell cycle-related genes is less pronounced in LUSC compared to LUAD.

Collectively, these results demonstrate that the level of BTNL9 protein sensitizes lung cancer cells to chemotherapy by promoting G0/G1 and G2/M cell cycle arrest through the downregulation of key cell cycle regulatory genes. These findings clearly highlight BTNL9 as a crucial factor affecting the efficacy of chemotherapy in lung cancer.

### Multifaceted impact of BTNL9 on the progression and prognosis of lung cancers

To assess the clinical relevance of BTNL9’s tumor-suppressive role, as observed in our in vitro and in vivo studies, we first performed immunohistochemistry (IHC) staining on paraffin-embedded tissues from normal and lung cancer patients. IHC staining revealed the presence of BTNL9 in the lung cancer cells of LUAD samples, with its expression exhibiting varying intensities – high, intermediate, and low (Fig. 8A). These findings are consistent with TCGA RNA-seq data, which show a broad spectrum of BTNL9 mRNA expression levels (Fig. 1B). Additionally, high-magnification images of normal lung tissues clearly demonstrated that BTNL9, expressed in pneumocytes, is localized in the nucleus (Fig. S6A).

**Fig. 8.**
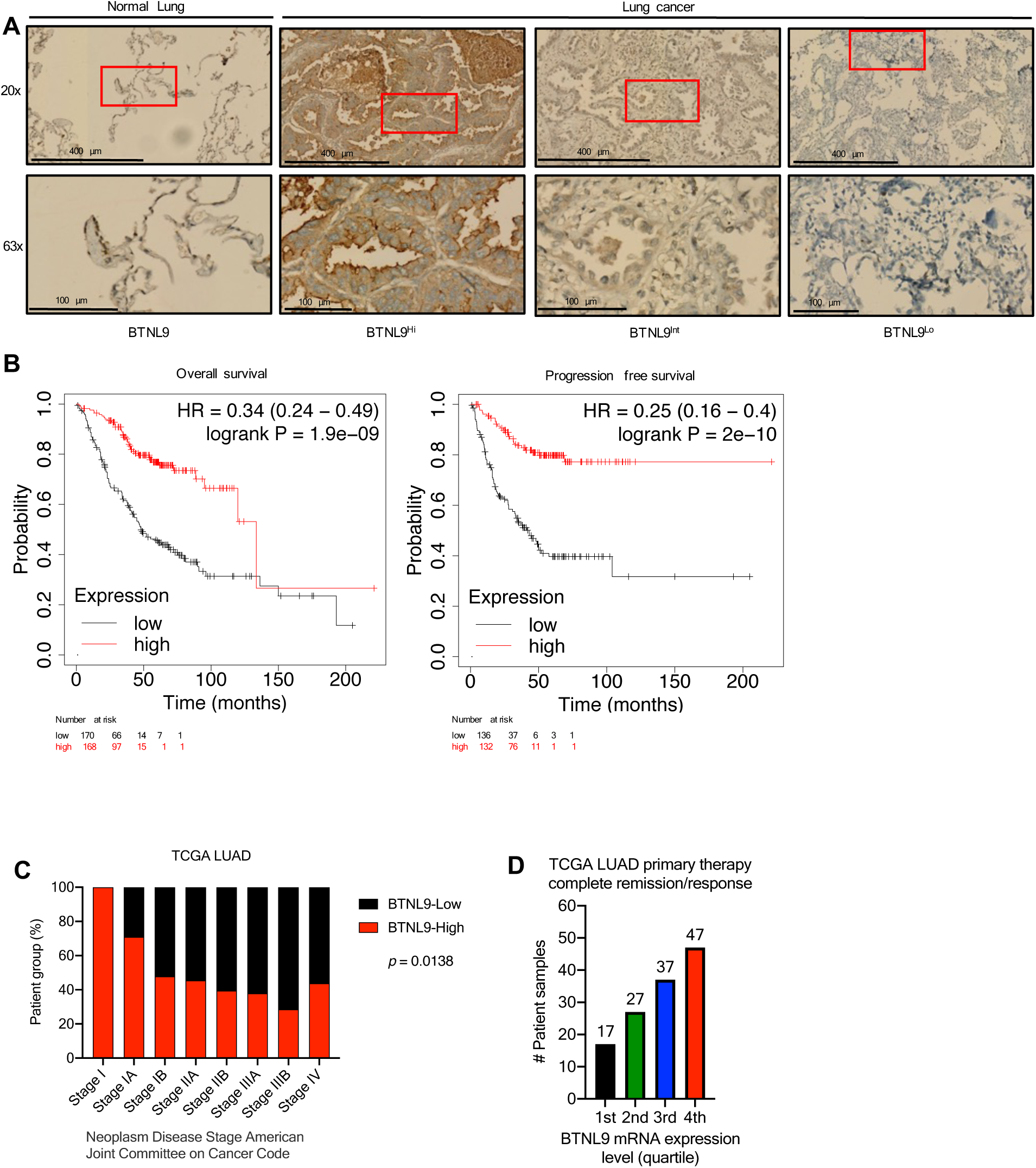
**Multifaceted impact of BTNL 9 on the progression and prognosis of lung cancers**. **A**, IHC staining on selected LUAD tumor and normal tissue samples from a tissue microarray, with images displayed at 20X and 63X magnification. The red box indicates an area of higher magnification. The BTNL9 stainings w ere categorized into high (BTNL9^Hi^), intermediate (BTNL9^I nt^), and low (BTNL9^Lo^) expression based on intensity. **B**, Kaplan–Meier curve analysis comparing overall survival and progression-free survival in LUAD patients within the first and fourth quartiles of BTNL9 expression. *p*-value was obtained from the log-tank test. **C**, Bar graph showing the distribution of LU AD patients with the first and fourth quartiles of BTNL9 expression according to tumor stages (as per America Joint Committee on Cancer (AJCC) Tumor Stage Code). Data was generated using TCGA LU AD dataset. **D**, Bar graph depicting the number of LUAD patients with complete remission/response follwing primary treatment, distributed across the four quartiles of BTNL9 expression.

Using the Kaplan–Meier plotter database, we analyzed data from 672 LUAD cases and 527 LUSC cases to generate Kaplan–Meier curves (30). We divided BTNL9 expression levels into four quartiles for detailed comparison. Comparisons of survival rates between the lowest (1st quartile, BTNL9-Low) and highest (4th quartile, BTNL9-High) levels of BTNL9 demonstrated that LUAD patients with higher BTNL9 expression showed improvements in both overall survival and progression-free survival (Fig. 8B). Conversely, no significant correlation between survival rates and BTNL9 expression level was observed for LUSC (Fig. S6B). In addition, we further examined the overall survival of patients with various other cancer types. Our findings revealed that patients with breast cancer, liver cancer, clear cell renal cell carcinoma, and pancreatic ductal adenocarcinoma also experienced significantly better overall survival rates. This suggests that higher BTNL9 expression not only benefits lung cancer patients but also those with a range of other cancers (Fig. S6C).

Next, we selected TCGA data for LUAD and LUSC from the cBioPortal database for further analysis. Patients were grouped based on BTNL9-Low and BTNL9-High expression levels and then sorted according to tumor stage (Fig. 8C and Fig. S6D). Notably, LUAD patients with BTNL9-High expression were predominantly found in the earlier stages of tumors, particularly stage 1 and 1A. Conversely, LUAD patients exhibiting BTNL9-Low expression were more frequently found in the advanced tumor stages, especially stage IIB onward. However, the trend was not significant in the LUSC. This analysis indicates that BTNL9 expression level could significantly impact the progression of cancer cells into certain tumor stages. Moreover, this influence appears to be context-dependent, varying among different cell subtypes. Next, we investigated whether BTNL9 expression levels affect treatment outcomes. Patients exhibiting BTNL9-High expression showed a higher incidence of complete remission compared to those with BTNL9-Low expression (Fig. 8D and Fig. S6E). This finding highlights the potential of BTNL9 as a prognostic marker in predicting therapeutic outcomes.

Finally, we investigated the potential roles of BTNL9 in immune cell infiltration into the tumor microenvironment (TME) using TISIDB, a web portal for tumor and immune system interaction. TCGA analysis uncovered a direct positive correlation between BTNL9 expression and the presence of infiltrated immune cells, including follicular helper T cells (Tfh), activated B cells, macrophages, effector CD8+ T cells, and natural killer (NK) cells, in both the LUAD and LUSC samples (Fig. S6F). Interestingly, this correlation appeared slightly more pronounced in the LUSC samples than in the LUAD samples. These findings suggest that BTNL9 may play a role in tumor immunity during the tumor progression. Overall, these analyses imply that BTNL9 impacts the progression and clinical outcome of lung cancer, even though its effect appears to vary between LUSC and LUAD.

## DISCUSSION

In this study, we demonstrated BTNL9 functions as a bona fide transcription factor with novel tumor suppressor activity in lung cancers. Subcellular fractionation analysis indicated that at least three distinct BTNL9 isoforms are expressed in lung cancer cell lines. The isoform with the highest MW was primarily localized in the nucleus, whereas the other two were found mostly in the cytoplasm. Despite the presence of the NLS in these potential isoforms, which typically directs proteins to the nucleus, the two smaller isoforms remain in cytoplasm. While it is possible that under certain condition, cytoplasmic BTNL9 could translocate to the nucleus, this finding also offers a different perspective, implying that BTNL9 might exert different functions depending on its localization (31, 32). It is reasonable to assume that BTNL9 localized in the nucleus modulates transcriptional activities. Investigating the specific roles of the cytoplasmic isoforms of BTNL9 also provide a promising research direction. Given that up to 10 BTNL9 transcript variants have been reported, future research should focus on determining which variants are translated into BTNL9 protein isoforms. Even though BTNL9 possesses only a single leucine heptad repeat and has a less conserved basic region compared to other bZIP family transcription factors, our ChIP-seq data revealed that BTNL9 not only binds to promoter regions either as a transcriptional enhancer or repressor, but also binds to intronic and intergenic regions; these are common features that are shared by other transcription factors (23, 24). In this study, 87% of the identified ChIP peaks were located within the intronic, and intergenic regions, and these results are suggestive of a diverse range of regulatory activities of BTNL9 in gene regulation. These may include functions as a trans-acting factor, participation in intron-mediated enhancement, (33, 34) or involvement in non-coding RNA regulation (35), all of which are potential avenues for future exploration of the regulatory role of BTNL9.

RNA-seq analysis further indicated that BTNL9 actively and selectively modulates transcription of genes associated with diverse biological pathways including cell cycle, p53-mediated transcription regulation, and DNA replication, all of which are key processes involved in tumorigenesis on the top pathways to demonstrate its novel tumor-suppressive regulation functions. Specifically, BTNL9 transcriptionally regulates the MCM family genes, licensing factors, and cell cycle genes, a finding implying its crucial roles in maintaining genomic integrity and cell cycle progression. Adding another layer of complexity, BTNL9 regulates a group of p53-associated genes in a p53-dependent manner, hinting at broader role for BTNL9 in cancer cell biology complementary to p53 signaling. Similar mechanisms have been previously reported, in which NF-Y, TFE3 and TFEB demonstrated p53-dependent tumor suppression regulation (36, 37). It is notable that the 12 genes selected for validation in our study are previously well-characterized and are among the key players in tumorigenesis. For instance, multiple MCM genes are recognized as sensitive markers in premalignant lung cells, underlining their substantial relevance to tumor progression (38–40). Additionally, some of the genes down-regulated by BTNL9, such as *CDKN1A*, *CDC25C*, *CDK1*, *E2F1*, *FOXM1*, *MCM2* and *MCM7*, are known targets for tumor inhibitors. This suggests a potential synergy between BTNL9’s regulatory action and current cancer therapies (41–46).

Through our xenograft model, we established BTNL9’s role as a tumor suppressor gene. This aligns with our expectations because BTNL9 transcriptionally and translationally downregulated genes implicated in cell proliferation and DNA replication, thereby suppressing tumorigenicity. Based on our comprehensive analysis of the TCGA datasets, we infer that the silencing or downregulation of BTNL9 may be associated with tumor progression and possibly result in less favorable clinical outcomes, including reduced survival rates. This trend is more pronounced in LUAD than in LUSC and implies potential variations in BTNL9 functions specific to different cancer subtypes. In the case of these findings, the observed difference could stem from the different underlying causes of adenocarcinoma and squamous cell carcinoma. Nonetheless, the significant association observed between BTNL9 expression levels and complete remission rates after primary treatment is indicative of the potential of BTNL9 as a predictive biomarker for treatment response in both LUAD and LUSC. Furthermore, higher survival rates with increased BTNL9 mRNA expression point to its value as a prognostic marker that can offer insights into lung cancer clinical outcomes.

The tumor immune microenvironment (TIME) orchestrates an intricate relationship between tumor cells and infiltrating immune cells. This dynamic interplay determines the balance between anti-tumor and pro-tumor immunity, which in turn influences both tumor progression and treatment efficacy (47–49). Our TCGA LUAD and LUSC analysis demonstrated a substantiate correlation between BTNL9 expression and prominent infiltration of anti-tumor immune cells, including cytotoxic T cells, NK cells, and B cells. This finding also implies that BTNL9 might play a role in determining the clinical prognosis and response to treatment. Further studies are required to determine how BTNL9 promotes the infiltration of immune cells into tumors.

To explore whether BTNL9 protein level affected treatment efficacy, etoposide and bortezomib were selected as model drug agents for this study due to their distinct mechanisms of action, both targeting critical pathways involved in cancer cell survival and proliferation. Etoposide, a topoisomerase II inhibitor, induces DNA damage and apoptosis (50), whereas bortezomib, a 26S proteasome inhibitor, disrupts protein degradation pathways, including the turnover of key cell cycle regulators (51). Our findings indicate that both drugs selectively sensitize cancer cell lines, though with different preferences. A549 cells exhibited broad sensitization across various concentrations when treated with bortezomib, whereas etoposide had a more pronounced effect at lower doses. In contrast, NCI-H460 cells showed only marginal sensitization to etoposide compared to bortezomib which produced great sensitization in BTNL9 overexpression cells. Overall, both drugs significantly suppressed key cell cycle regulators, particularly in cancer cells with high level of BTNL9. One of these regulators, FOXM1, is a transcription factor that plays a pivotal role in cell cycle progression, especially during the G1/S and G2/M transition and mitotic progression (52) (53, 54). Its activation is also involved in the phosphorylation of CDK1, CDK2, and Cyclin B1 (*CCNB1*) and Aurora kinase B (*AURKB*)(55, 56). A negative correlation was found between BTNL9 and these key cell cycle regulators in LUAD TCGA data further supports the role of BTNL9 in their suppression. These findings suggest that BTNL9 expression levels in cancer cells play a crucial role in determining treatment efficacy. Based on this novel discovery, we propose that BTNL9 could serve as an important biomarker for guiding treatment selection.

## Conclusion

In summary, we have successfully identified BTNL9’s roles as a novel transcription factor with tumor-suppressive activities and delved into its unique features that could provide insights in the intricacies of cancer biology (Fig. 9). With this fresh perspective, BTNL9 might become a primary focus for future cancer therapeutic developments. There are still many unanswered questions regarding the biology of BTNL9. For example, the enigmatic suppression of BTNL9 mRNA expression in cancer, despite its low mutation rate, suggests that an unidentified mechanism for silencing its expression is at play. Future studies need to focus on the regulation of BTNL9 expression in cancers, as understanding its regulation could provide valuable insights for future therapeutic approaches.

**Fig. 9.**
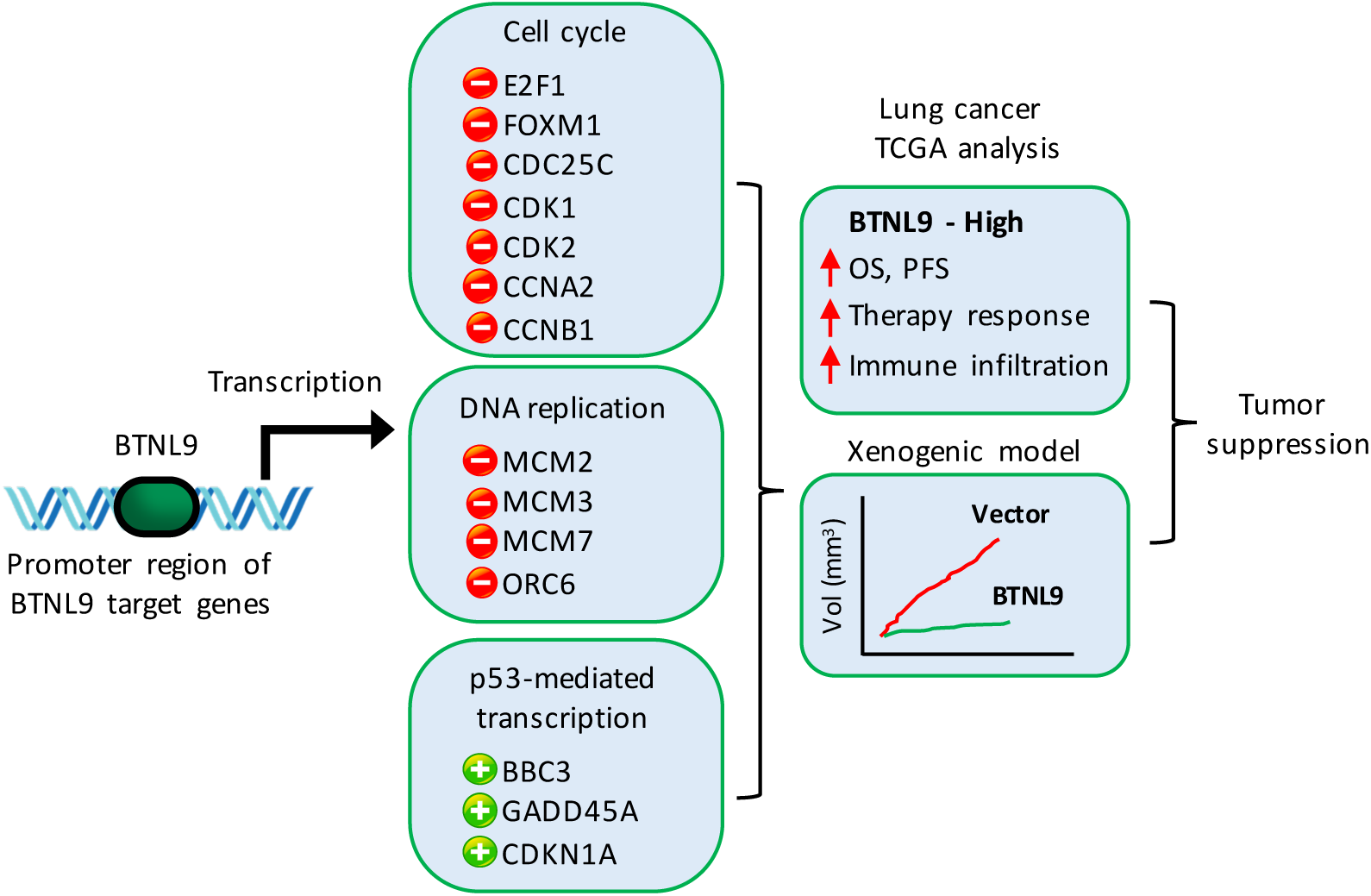
Schematic illustration of BTN L9’s function as a transcription factor with tumor suppressive activity. BTN L9 binds to promoter regions to regulate the transcription of target genes that are involved in cell cycle, DNA replication, and p53-mediated transcription. Higher BTNL9 expression levels are positively correlated with overall survival, progression-free survival, treatment outcomes, and immune infiltration. Xenograft mouse model provides strong evidence of BTN L9’s tumor-suppressive activity, demonstrating significantly reduced tumor growth at higher BTNL9 levels.

## DATA AND MATERIALS AVAILABILITY

All data needed to evaluate the conclusions in the paper are present in the paper and/or the Supplementary Materials. ChIP-seq and RNA-seq data have been deposited into the NCBI’s Short Read Archive (RRID:SCR_004801; BioProject ID PRJNA1011818) with Series GSE242317 (ChIP-seq) and Series GSE242319 (RNA-seq).

## ACKNOWLEDGMENTS

Funding: This research was supported by Basic Science Research Program through the National Research Foundation of Korea (NRF) funded by the Ministry of Education (2021R1I1A30489331361582062860103).

## AUTHOR CONTRIBUTIONS

PY, WLN designed and performed most of the experiment, data analysis and drafted the manuscript. HJC, MSK performed all the pathological works. IC, WLN supervised and finalized the manuscript. All authors read and approved the manuscript submission.

