## Supplemental figures and tables for "Novel transcription factor BTNL9 enhances tumor suppression and drug sensitivity in non-small cell lung cancer through cell cycle regulation"

### Supplementary Figures

#### Figure S1

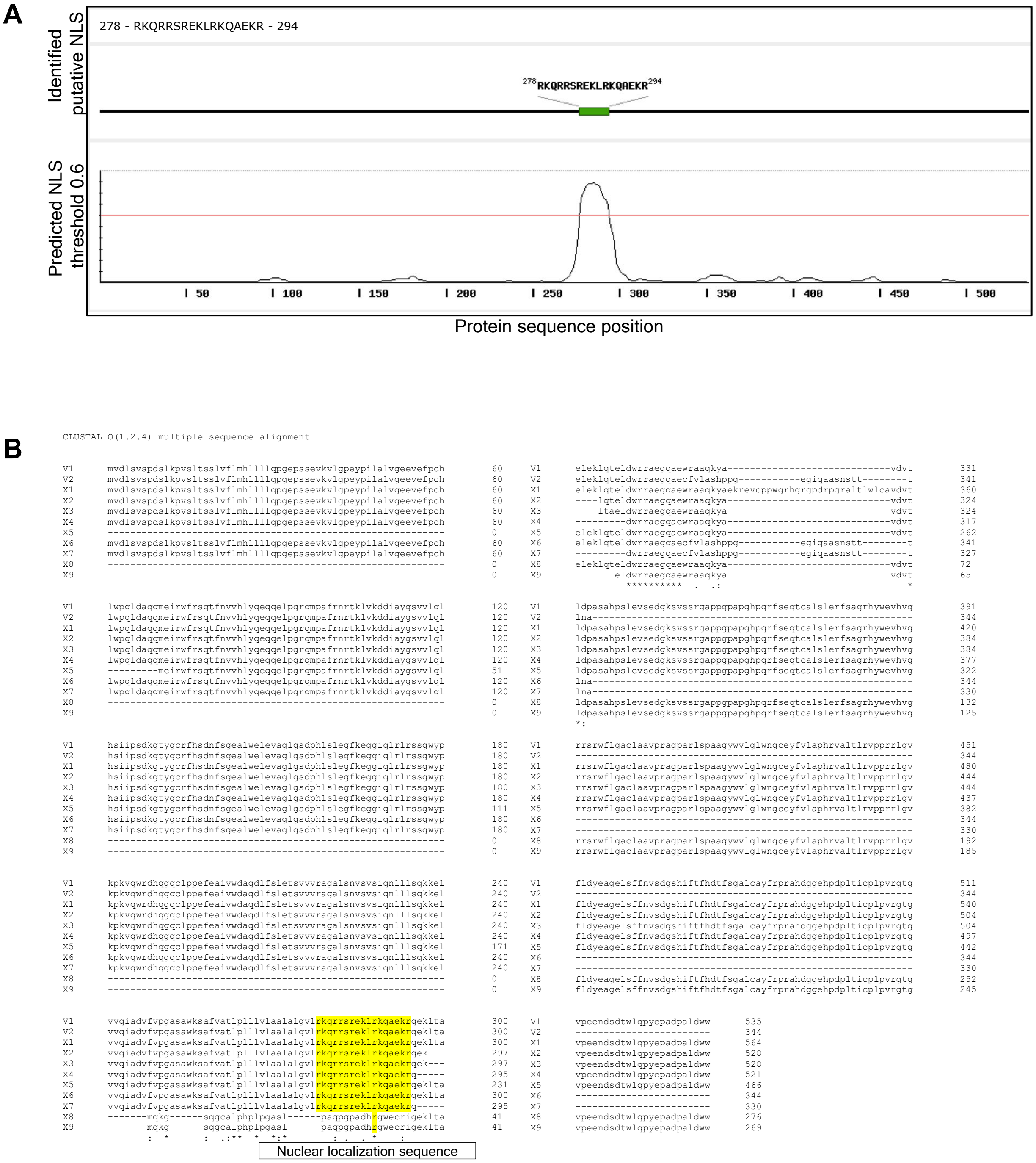


**Fig. S1 Nuclear localization sequence analysis.**

**A,** Full-length protein sequence of BTNL9 was analyzed using NLStradamus, a program to predict potential NLS domain. Short sequence RKQRRSREKLRKQAEKR was identified as NLS, with threshold score above 0.6. **B**, Protein sequence alignment of all available BTNL9 variants from NCBI revealed a conserved NLS domain except for variant X8 and X9.

**Figure S2**

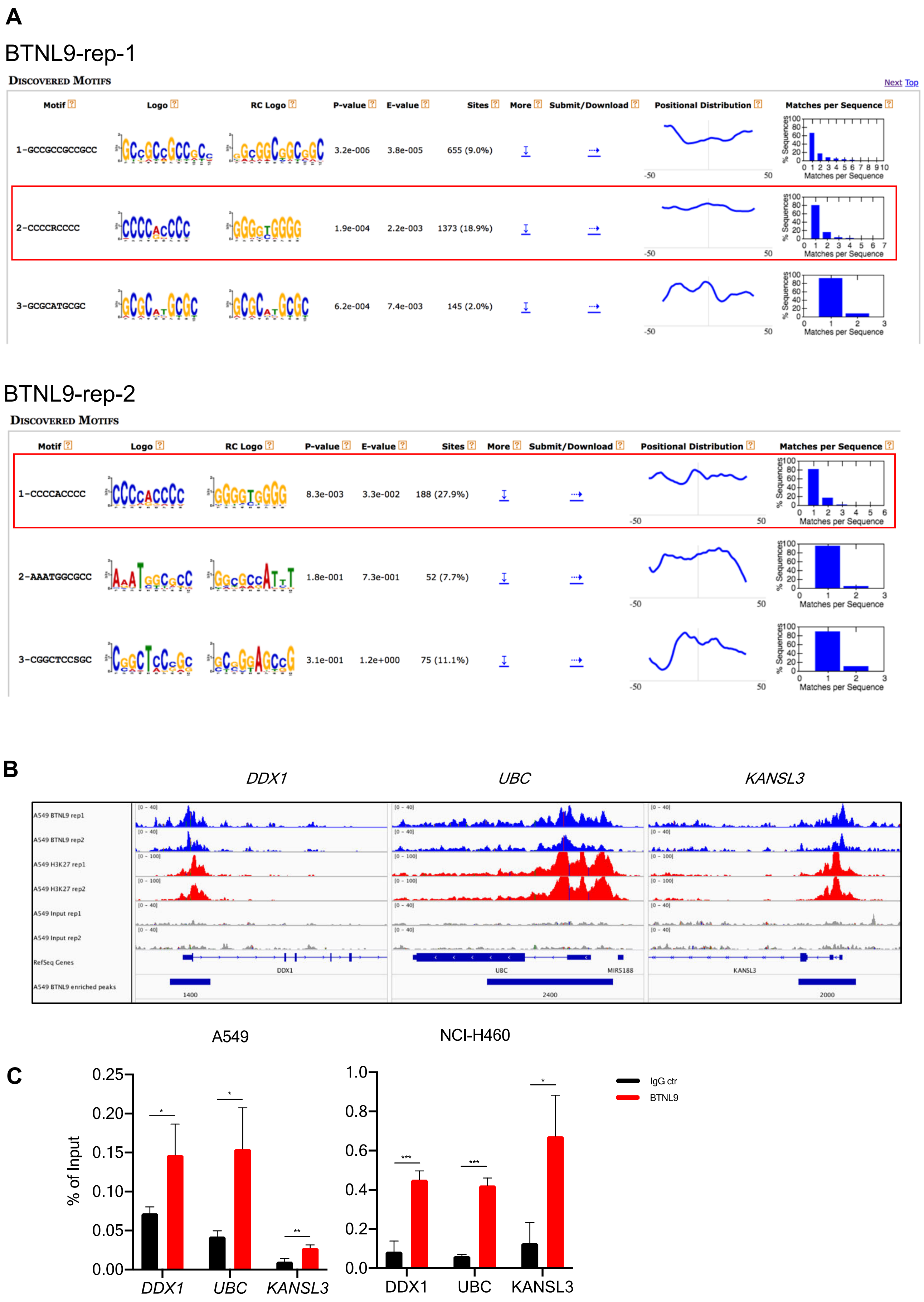
A
BTNL9-rep-1
Discovered Motifs
Next Top
Motif | Logo | RC Logo | P-value | E-value | Sites | More | Submit/Download | Positional Distribution | Matches per Sequence
1-GCCGCCGCCGCC | 3.2e-006 | 3.8e-005 | 655 (9.0%)
2-CCCCRCCCC | 1.9e-004 | 2.2e-003 | 1373 (18.9%)
3-GCGCATGCGC | 6.2e-004 | 7.4e-003 | 145 (2.0%)
BTNL9-rep-2
Discovered Motifs
1-CCCCACCCC | 8.3e-003 | 3.3e-002 | 188 (27.9%)
2-AAATGGCGCC | 1.8e-001 | 7.3e-001 | 52 (7.7%)
3-CGGCTCCSGC | 3.1e-001 | 1.2e+000 | 75 (11.1%)
% Sequences
Matches per Sequence
B
DDX1
UBC
KANSL3
A549 BTNL9 rep1
A549 BTNL9 rep2
A549 H3K27 rep1
A549 H3K27 rep2
A549 Input rep1
A549 Input rep2
RefSeq Genes
A549 BTNL9 enriched peaks
MIR5188
1400
2400
2000
A549
NCI-H460
C
% of Input
DDX1
UBC
KANSL3
IgG ctr
BTNL9

#### Figure S2

D

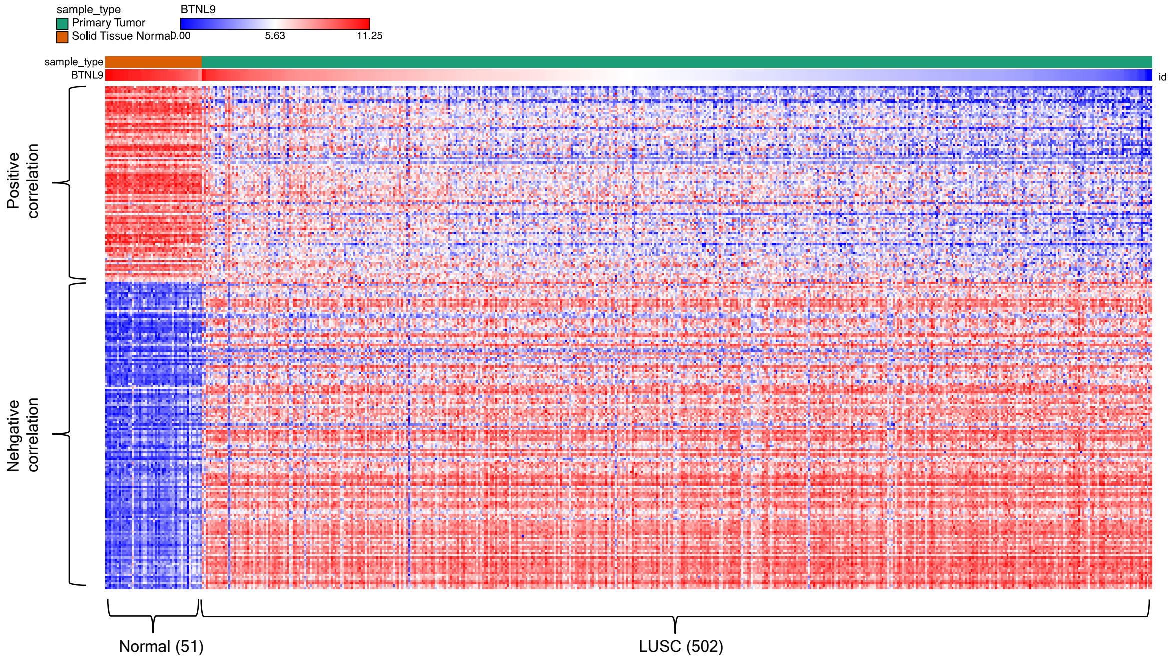


**Fig. S2 Potential target genes and transcription factors of BTNL9 and its binding regions.**

**A**, MEME-ChIP DNA motif analyses of both BTNL9 ChIP-seq replicates revealed a potential DNA binding conserved motif sequence. The red box highlights the consistent presence of the same motif CCCCRCCCC in both sets of ChIP-seq data. **B**, . Integrative Genomic Viewer (IGV) pot displays ChIP-seq peak of BTNL9, H3K27ac, and input control at the loci of the random selected genes (*DDX1, UBC,* and *KANSL3*) for ChIP-qPCR validation. **C**, Bar graphs representing the ChIP-qPCR validation of chromatin enrichment for 3 genes (*DDX1*, *UBC* and *KANSL3*), as identified in ChIP-seq data. Results were presented as % of input derived from delta CT between ChIP and 100% input. Significance was determined by a two-tailed Student's *t*-test: * $p < 0.05$ and ** $p < 0.01$. **D**, Heatmap illustrating expression profiles of 46 genes significantly correlated to BTNL9 mRNA levels from TCGA LUSC dataset (Spearman's correlation coefficient $r \leq$ - 0.45 or $r \geq 0.45$).

#### Figure S3

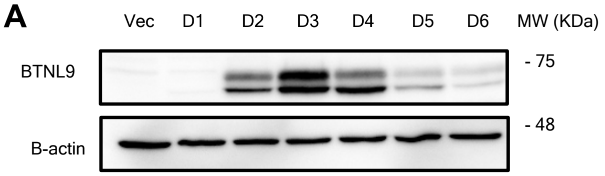


**Fig. S3 BTNL9 expression profiles.**

**A**, Western blot showing the temporal expression pattern of BTNL9 in A549 that were infected with either a empty vector control or BTNL9 lentivirus and harvested from Day 1 to Day 6, using β-actin as a loading control.

### Figure S4

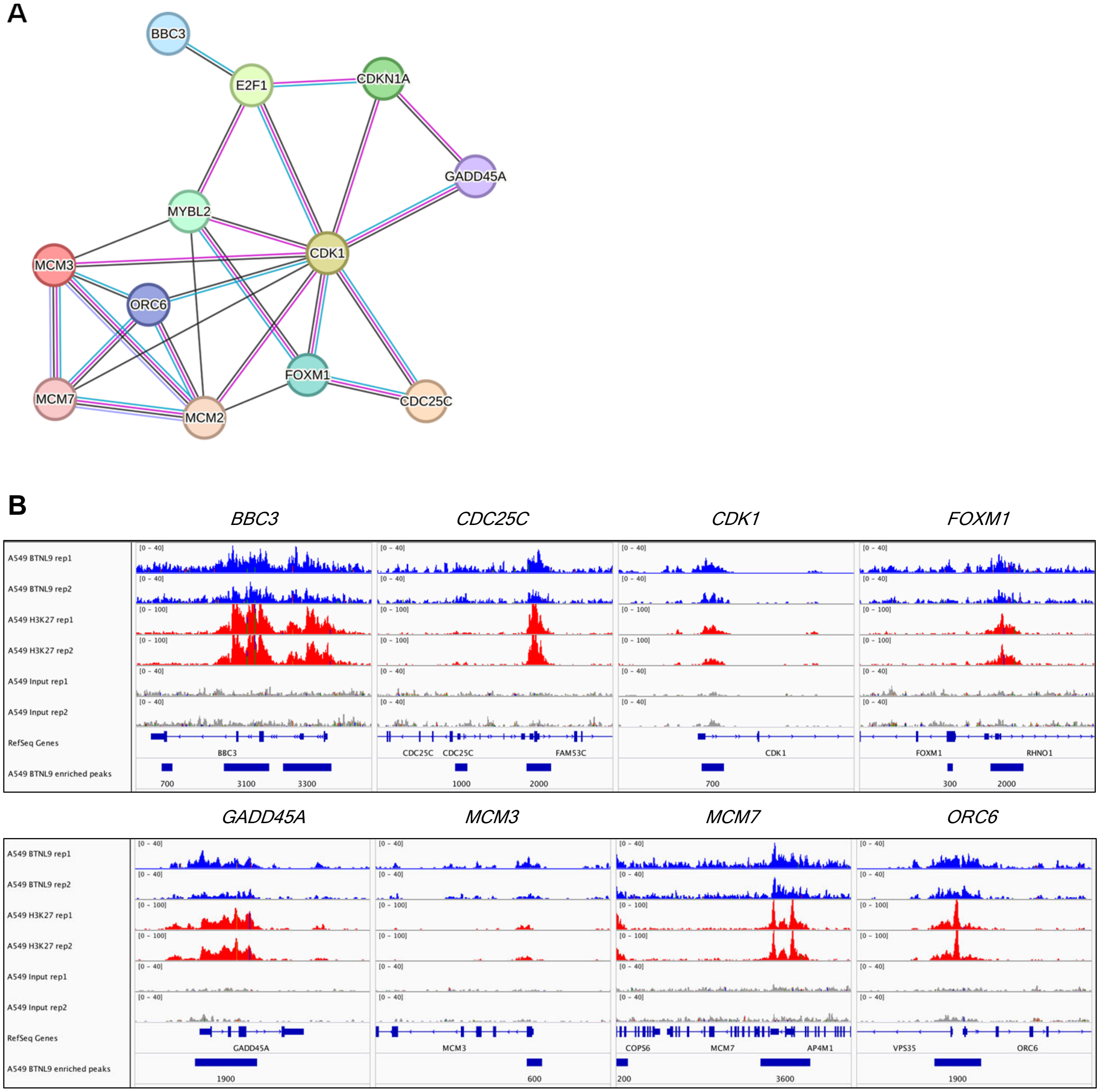
A
BBC3
E2F1
CDKN1A
GADD45A
MYBL2
CDK1
MCM3
ORC6
FOXM1
MCM7
MCM2
CDC25C
B
BBC3
CDC25C
CDK1
FOXM1
GADD45A
MCM3
MCM7
ORC6
A549 BTNL9 rep1
A549 BTNL9 rep2
A549 H3K27 rep1
A549 H3K27 rep2
A549 Input rep1
A549 Input rep2
RefSeq Genes
A549 BTNL9 enriched peaks
[0 - 40]
[0 - 100]
BBC3
700
3100
3300
CDC25C
CDC25C
FAM53C
1000
2000
CDK1
700
FOXM1
RHNO1
300
2000
GADD45A
1900
MCM3
600
COPS6
MCM7
AP4M1
200
3600
VPS35
ORC6
1900

### Figure S4

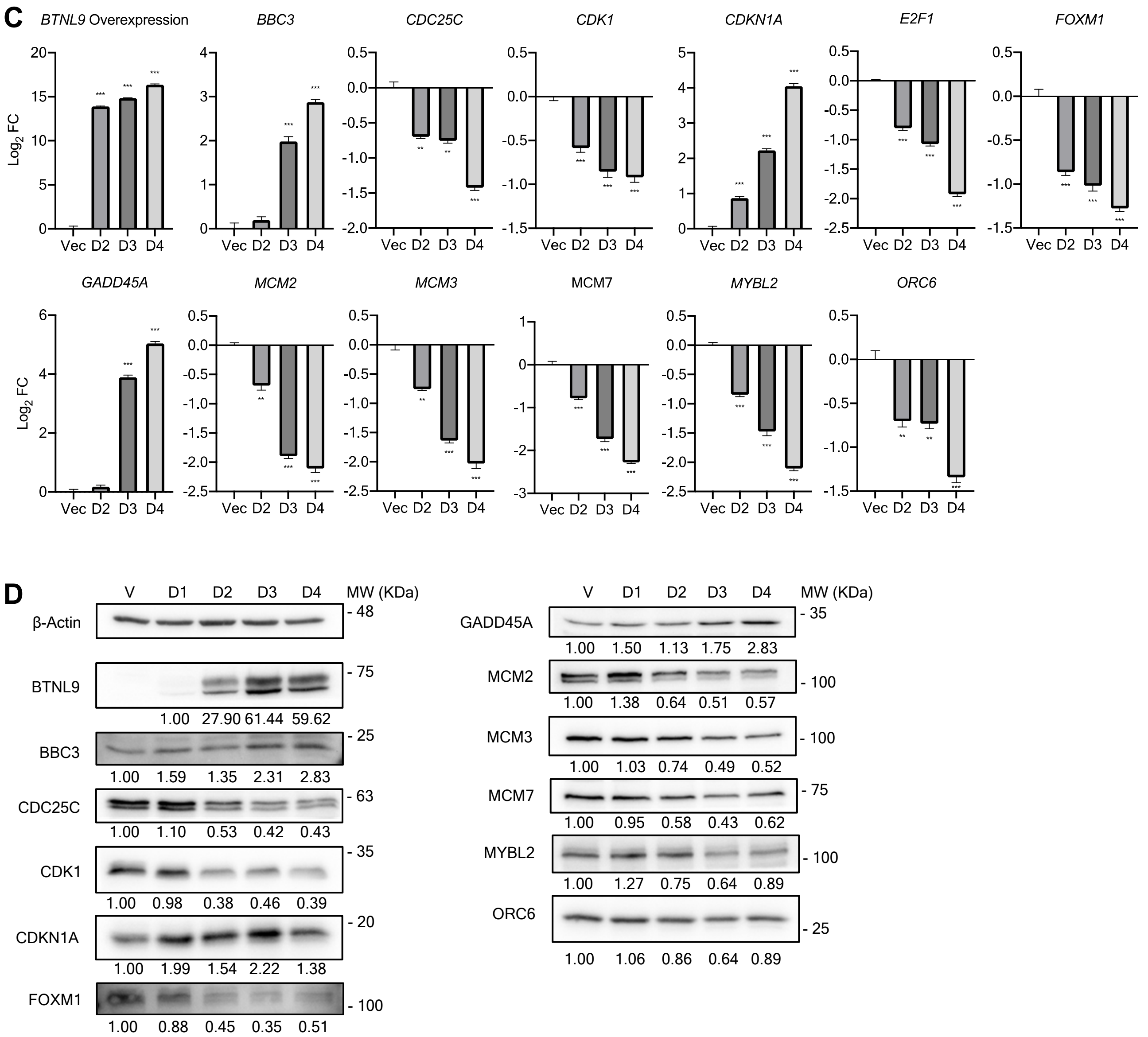
C
BTNL9 Overexpression
BBC3
CDC25C
CDK1
CDKN1A
E2F1
FOXM1
GADD45A
MCM2
MCM3
MCM7
MYBL2
ORC6
Log2 FC
Vec D2 D3 D4
D
V D1 D2 D3 D4 MW (KDa)
β-Actin - 48
BTNL9 - 75
1.00 27.90 61.44 59.62
BBC3 - 25
1.00 1.59 1.35 2.31 2.83
CDC25C - 63
1.00 1.10 0.53 0.42 0.43
CDK1 - 35
1.00 0.98 0.38 0.46 0.39
CDKN1A - 20
1.00 1.99 1.54 2.22 1.38
FOXM1 - 100
1.00 0.88 0.45 0.35 0.51
GADD45A - 35
1.00 1.50 1.13 1.75 2.83
MCM2 - 100
1.00 1.38 0.64 0.51 0.57
MCM3 - 100
1.00 1.03 0.74 0.49 0.52
MCM7 - 75
1.00 0.95 0.58 0.43 0.62
MYBL2 - 100
1.00 1.27 0.75 0.64 0.89
ORC6 - 25
1.00 1.06 0.86 0.64 0.89

**Figure S4**

E

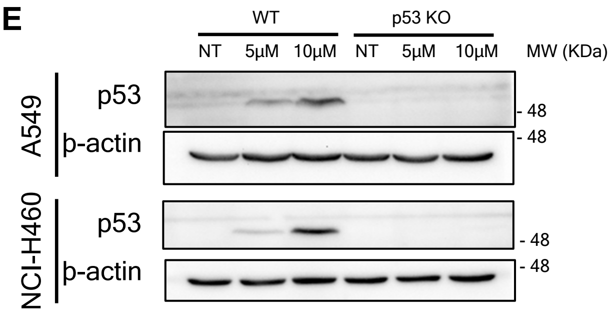
WT
p53 KO
NT
5μM
10μM
NT
5μM
10μM
MW (KDa)
A549
p53
β-actin
NCI-H460
p53
β-actin
- 48
- 48
- 48
- 48

F

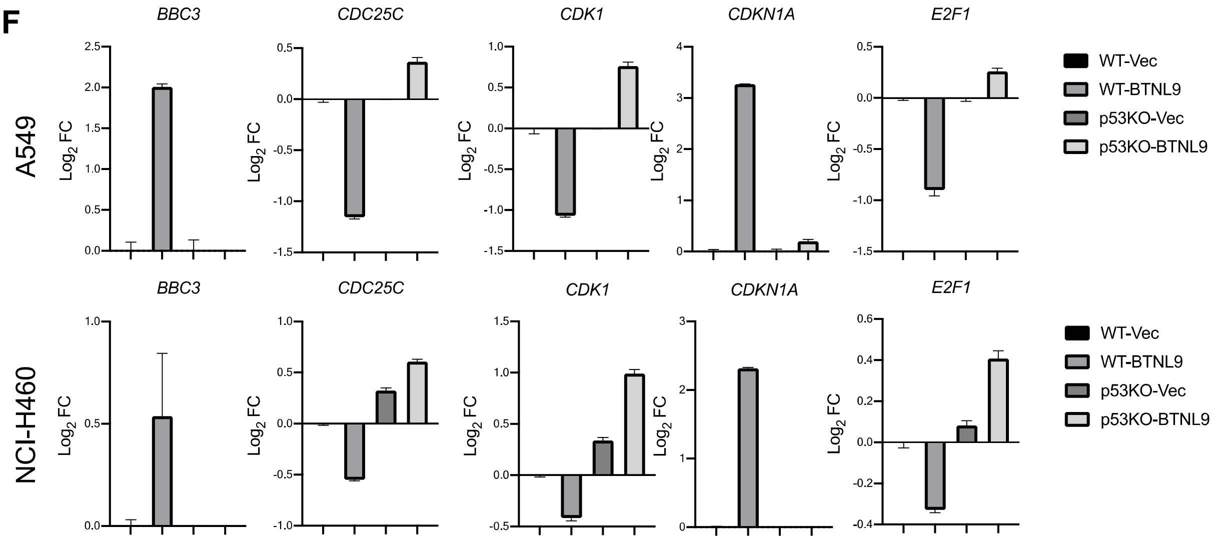
BBC3
CDC25C
CDK1
CDKN1A
E2F1
A549
NCI-H460
Log2 FC
WT-Vec
WT-BTNL9
p53KO-Vec
p53KO-BTNL9

**Fig. S4 Analysis of BTNL9 RNA-seq data obtained in NCI-H460 cell line**

**A**, String analysis showing interconnection between the selected 12 genes from three clusters identified in RNA-seq data. **B**, Integrative Genomics Viewer (IGV) plot showcasing ChIP-seq peaks of BTNL9, H3K27ac, and input controls at the loci of *MCM7* and *BBC3*. **C**, Bar graphs representing qRT-PCR analyses in NCI-H460 cells overexpressing either an empty vector control or BTNL9, with samples harvested on the indicated day. All values were calculated as mean Log2 FC ±SD. Significance was determined by a two-tailed Student's *t*-test: * $p < 0.05$, ** $p < 0.01$ and *** $p < 0.001$. All experiments were conducted in triplicates. Primers for qRT-PCR of tested genes were listed in Table S8. **D**, Representative western blot for selected genes in NCI-H460 cells overexpressing either an empty vector control or BTNL9, with samples harvested on the indicated day. β-actin was used as a loading control. Signal intensities were quantified with iBright Analysis software and semi-quantitative protein expression levels were calculated by normalizing to β-actin. E2F1 was not included in the western blot results as its signal was underdetectable. **E**, Western blot analysis of p53 expression in wild-type (WT) and p53 knockout (p53 KO) A549 and NCI-H460 cell lines. Cells were treated with Nutlin-3 at 5 μM and 10 μM for 24 hours. þ-actin was used as a housekeeping control. **F**, mRNA expression levels of BTNL9 downstream target genes (*BBC3*, *CDC25C*, *CDK1*, *CDKN1A*, and *E2F1*) were assessed in WT and p53 KO A549 and NCI-H460 cells using RT-qPCR. Primers for qRT-PCR were listed in Table S8. For **C** and **F** data represents the mean of biological triplicates and tested by Student's *t* test (unpaired two-tailed). * $p < 0.05$, ** $p < 0.01$ and *** $p < 0.001$.

### Figure S5

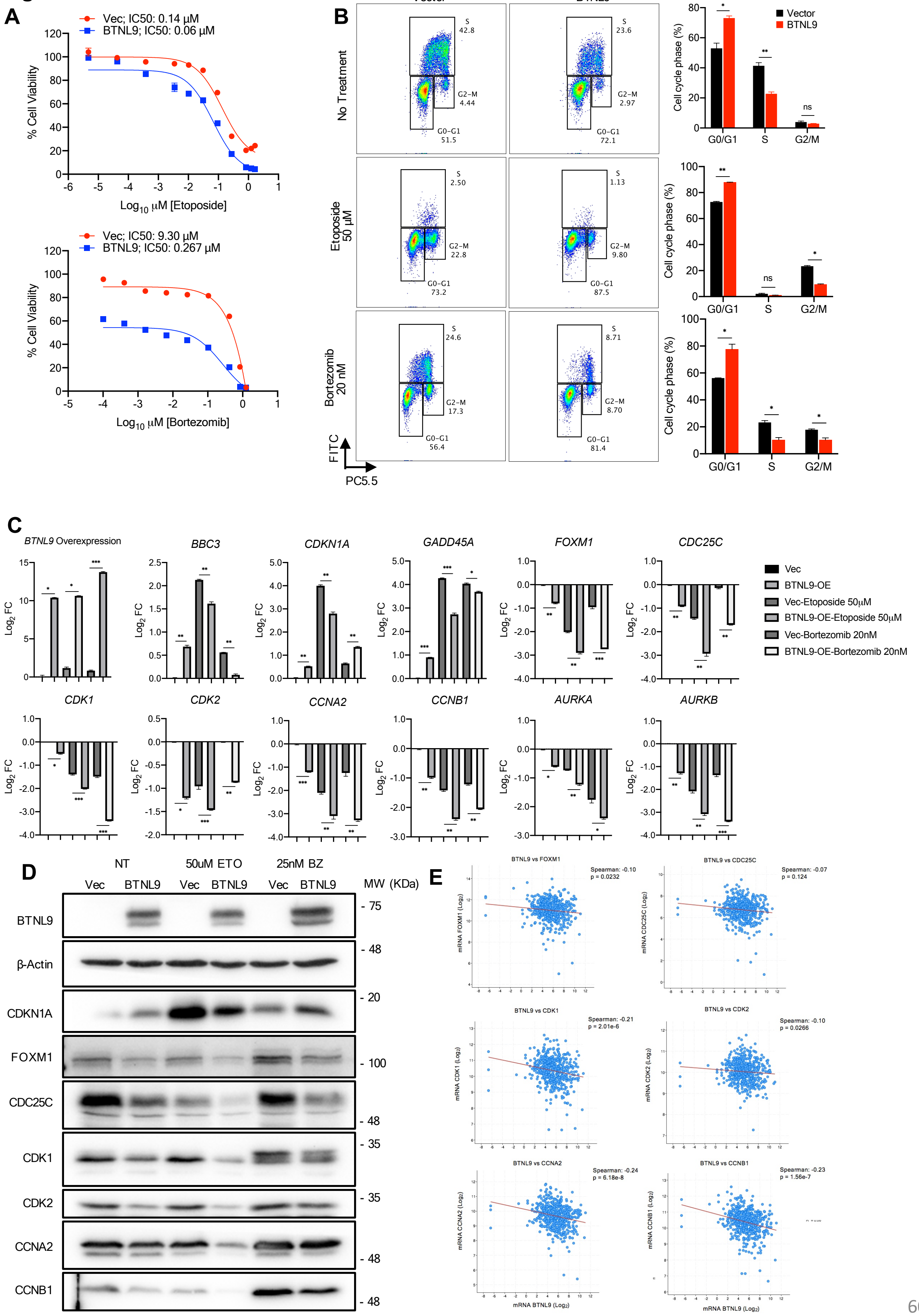

A
Vec; IC50: 0.14 µM
BTNL9; IC50: 0.06 µM
% Cell Viability
Log10 µM [Etoposide]
Vec; IC50: 9.30 µM
BTNL9; IC50: 0.267 µM
Log10 µM [Bortezomib]
B
Vector
BTNL9
No Treatment
Etoposide 50 µM
Bortezomib 20 nM
FITC
PC5.5
Cell cycle phase (%)
G0/G1
S
G2/M
C
BTNL9 Overexpression
BBC3
CDKN1A
GADD45A
FOXM1
CDC25C
CDK1
CDK2
CCNA2
CCNB1
AURKA
AURKB
Log2 FC
Vec
BTNL9-OE
Vec-Etoposide 50µM
BTNL9-OE-Etoposide 50µM
Vec-Bortezomib 20nM
BTNL9-OE-Bortezomib 20nM
D
NT
50uM ETO
25nM BZ
Vec
BTNL9
MW (KDa)
BTNL9
β-Actin
CDKN1A
FOXM1
CDC25C
CDK1
CDK2
CCNA2
CCNB1
E
BTNL9 vs FOXM1
BTNL9 vs CDC25C
BTNL9 vs CDK1
BTNL9 vs CDK2
BTNL9 vs CCNA2
BTNL9 vs CCNB1
mRNA BTNL9 (Log2)

**Fig. S5 BTNL9 enhances drug sensitivity to etoposide and bortezomib in lung cancer cells by regulating cell cycle pathways in NCI-H460.**

**A**, Dose-response curves from cytotoxicity assays of etoposide and bortezomib treatments for 48 hr in NCI-H460 cells overexpressing BTNL9 or vector control was used to estimate for IC50 value. **B**, Representative scatter plots for the cell cycle assay, performed using BrdU staining on vector control and BTNL9-overexpressing in NCI-H460 cells. Cell populations representing G0/G1 phase, S phase and G2/M phase were gated. Bar graphs representing the compilation of individual experimental results with mean ± SD. **C**, Bar graphs representing RT-qPCR analyses in NCI-H460 cells overexpressing either empty vector control or BTNL9. All values were calculated as mean $Log_2$ FC ±SD. Significance was determined by a two-tailed Student's *t*-test: * $p < 0.05$, ** $p < 0.01$ and *** $p < 0.001$. All experiments were conducted in biological triplicates. Primers for qRT-PCR of tested genes were listed in Table S8. **D**, Representative western blot of selected key cell cycle markers in NCI-H460 cells overexpressing either empty vector control or BTNL9. β-actin was used as a loading control. **E**, Correlation analysis between BTNL9 mRNA levels and the expression of cell cycle-related genes (*FOXM1*, *CDC25C*, *CDK1*, *CDK2*, *CCNA2*, and *CCNB1*) in LUSC patient samples derived from TCGA dataset. Spearman correlation coefficients and p-values are shown, part of the cell cycle markers indicating a negative correlation with BTNL9 expression. For **B**, **C**, and **D**, cells were either with no treatment (NT), or treated with 50 μM etoposide or 20 nM bortezomib for 18 hr. For **B** and **C** data represents the mean of biological triplicates and tested by Student's *t* test (unpaired two-tailed). * $p < 0.05$, ** $p < 0.01$ and *** $p < 0.001$.

### Figure S6

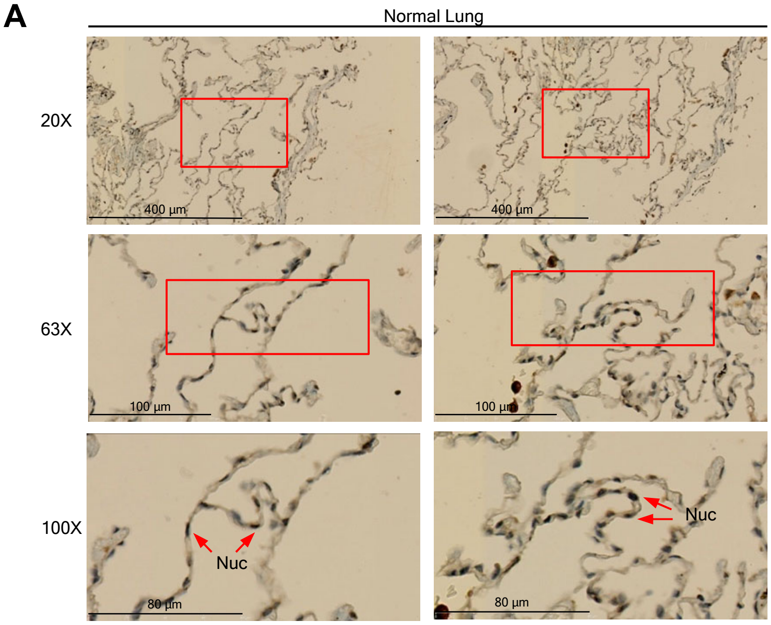
A
Normal Lung
20X
63X
100X
400 µm
400 µm
100 µm
100 µm
80 µm
80 µm
Nuc
Nuc

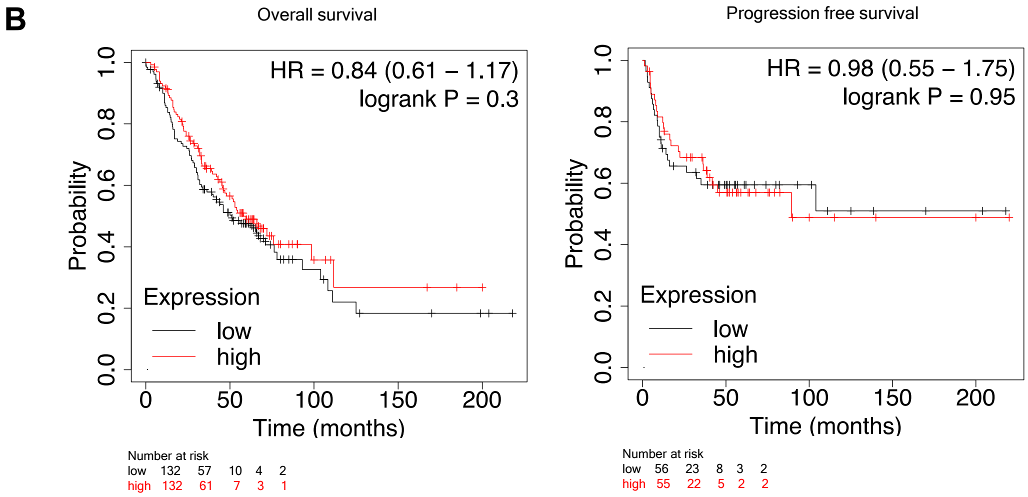
B
Overall survival
HR = 0.84 (0.61 – 1.17)
logrank P = 0.3
Probability
Time (months)
Expression
low
high
Number at risk
low 132 57 10 4 2
high 132 61 7 3 1
Progression free survival
HR = 0.98 (0.55 – 1.75)
logrank P = 0.95
Probability
Time (months)
Expression
low
high
Number at risk
low 56 23 8 3 2
high 55 22 5 2 2

### Figure S6

**C**

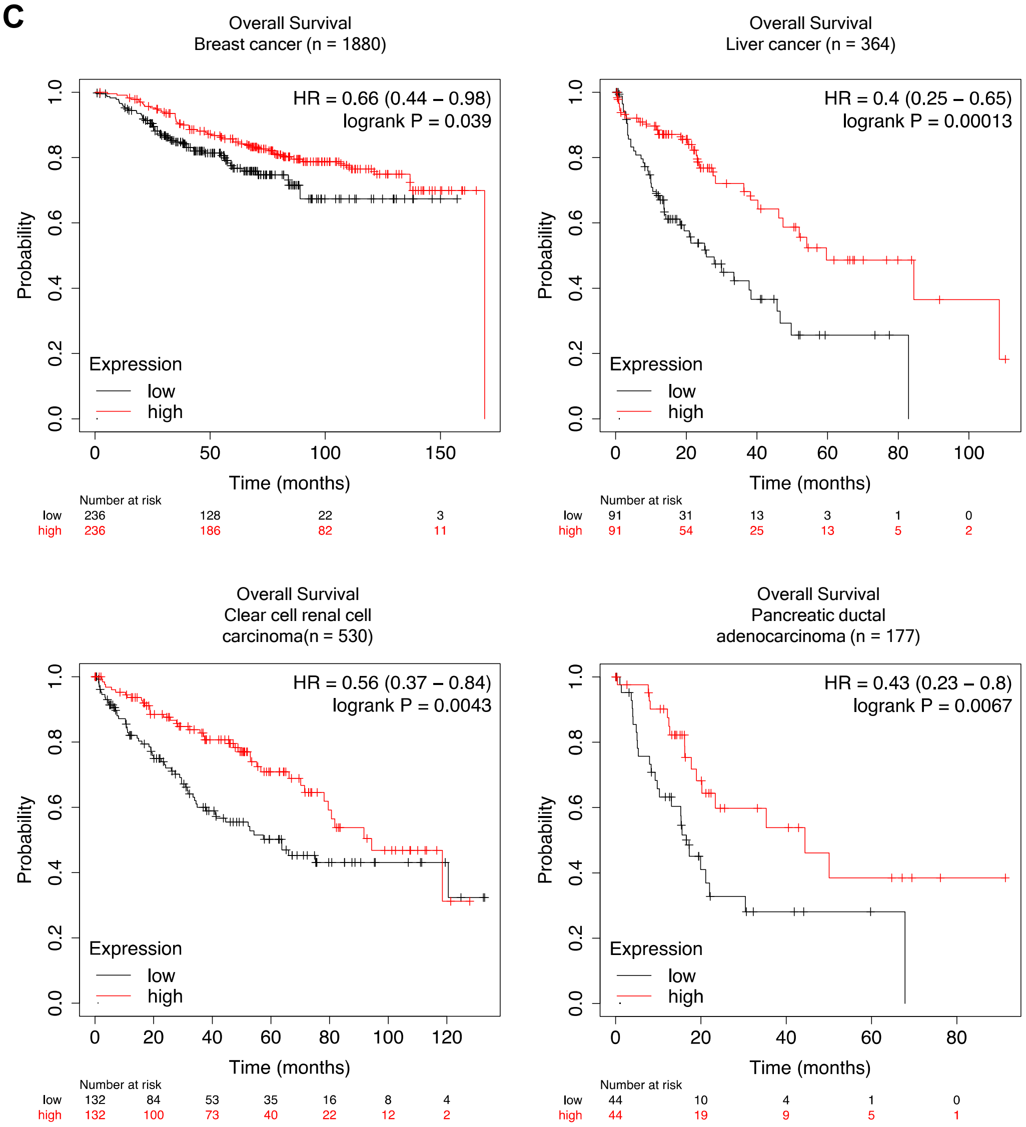
Overall Survival
Breast cancer (n = 1880)
HR = 0.66 (0.44 – 0.98)
logrank P = 0.039
Probability
Expression
low
high
Time (months)
Number at risk
low 236 128 22 3
high 236 186 82 11
Overall Survival
Liver cancer (n = 364)
HR = 0.4 (0.25 – 0.65)
logrank P = 0.00013
Probability
Expression
low
high
Time (months)
Number at risk
low 91 31 13 3 1 0
high 91 54 25 13 5 2
Overall Survival
Clear cell renal cell
carcinoma(n = 530)
HR = 0.56 (0.37 – 0.84)
logrank P = 0.0043
Probability
Expression
low
high
Time (months)
Number at risk
low 132 84 53 35 16 8 4
high 132 100 73 40 22 12 2
Overall Survival
Pancreatic ductal
adenocarcinoma (n = 177)
HR = 0.43 (0.23 – 0.8)
logrank P = 0.0067
Probability
Expression
low
high
Time (months)
Number at risk
low 44 10 4 1 0
high 44 19 9 5 1

### Figure S6

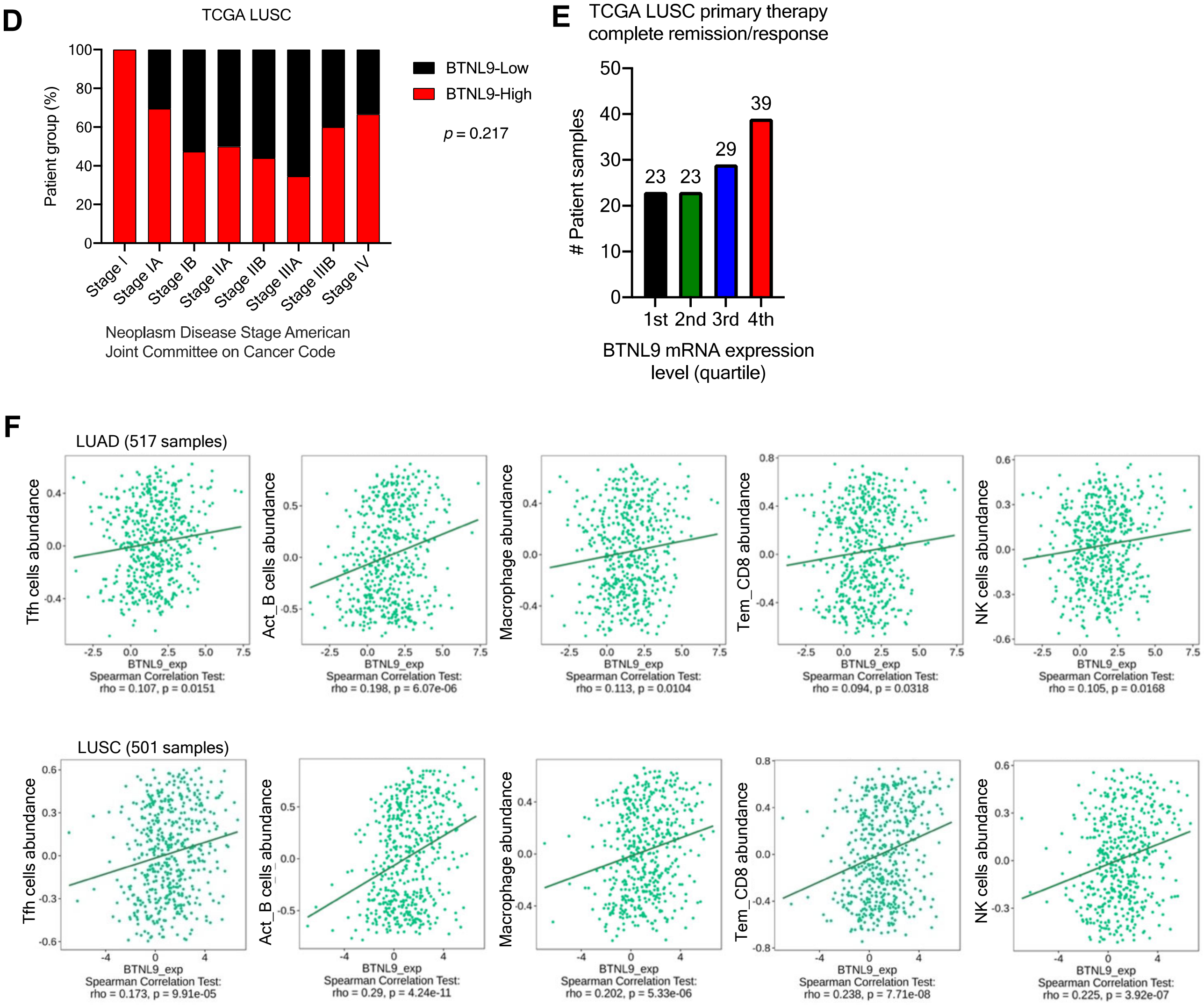
D
TCGA LUSC
Patient group (%)
100
80
60
40
20
0
Stage I
Stage IA
Stage IB
Stage IIA
Stage IIB
Stage IIIA
Stage IIIB
Stage IV
Neoplasm Disease Stage American Joint Committee on Cancer Code
BTNL9-Low
BTNL9-High
p = 0.217
E
TCGA LUSC primary therapy complete remission/response
### Patient samples
50
40
30
20
10
0
23
23
29
39
1st
2nd
3rd
4th
BTNL9 mRNA expression level (quartile)
F
LUAD (517 samples)
Tfh cells abundance
Act_B cells abundance
Macrophage abundance
Tem_CD8 abundance
NK cells abundance
BTNL9_exp
Spearman Correlation Test:
rho = 0.107, p = 0.0151
rho = 0.198, p = 6.07e-06
rho = 0.113, p = 0.0104
rho = 0.094, p = 0.0318
rho = 0.105, p = 0.0168
LUSC (501 samples)
rho = 0.173, p = 9.91e-05
rho = 0.29, p = 4.24e-11
rho = 0.202, p = 5.33e-06
rho = 0.238, p = 7.71e-08
rho = 0.225, p = 3.92e-07

**Fig. S6 Multifaceted impact of BTNL9 on the progression and prognosis of lung cancer.**

**A**, IHC staining on normal lung tissue samples from a tissue microarray. Images were captured at 20X and 63X magnifications, with areas inside of box further magnified to X100. Red arrows highlight pneumocytes exhibiting nuclear-localized BTNL9 expression. **B**, Kaplan–Meier curve analysis comparing overall survival and progression-free survival in LUSC patients within the first and fourth quartiles of BTNL9 expression. *p*-value was obtained from the log-rank test. **C**, Kaplan–Meier curve analysis comparing overall survival in cancers other than lung cancer, specifically breast cancer, liver cancer, clear cell renal cell carcinoma, and pancreatic ductal adenocarcinoma, within the first and fourth quartiles of BTNL9 expression. *p*-value was determined by the log-rank test. **D**, Bar graph showing the distribution of LUSC patients with the within the first and fourth quartiles quartiles of BTNL9 expression according to tumor stages (as per America Joint Committee on Cancer (AJCC) Tumor Stage Code). Data was generated using TCGA LUAD dataset. **E**, Bar graph depicting the number of LUSC patients with complete remission/response following primary treatment, distributed across the four quartiles of BTNL9 expression**F**, Correlation analysis of BTNL9 gene expression with immune cell infiltration based on TCGA LUAD and LUSC dataset. A positive correlation with *rho* > 0 and a negative correlation with *rho* < 0. Significant data points are represented by $p < 0.05$. Five immune cell populations were examined: Follicular helper T cells (Tfh), activated B cells, macrophages, effector CD8+ T cell, and natural killer (NK) cells.

**Table S1**
A549 ChIP-seq peak call data for BTNL9 revealed a list of 9,707 genes, identified through stringent criteria with $p < 1.0E\text{-}5$ (MACS recommendations), and a false discovery rate (FDR) of $q < 0.005$. (**This is a data file in Excel spreadsheet.**)

**Table S2**
Genomic Regions Enrichment of Annotations Tool (GREAT) was used to annotate ChIP peaks to the nearest genes within ±1 kb of the transcription start site (TSS), identifying 9,707 genes associated with BTNL9 binding sites. (**This is a data file in Excel spreadsheet.**)

**Table S3**
Summary of genes identified from A549 ChIP-seq data with significant clinical relevance to BTNL9 expression in lung cancer patients. (Spearman's Correlation coefficient $r \leq -0.45$ or $r \geq 0.45$). (**This is a data file in Excel spreadsheet.**)

**Table S4**
BTNL9 binding site was identified in 594 gene encoded transcription factor. (**This is a data file in Excel spreadsheet.**)

**Table S5**
List of 444 genes from A549 RNA-seq genes that fit the criteria of $Log_2$ FC ± 0.5 and FDR $q < 0.1$. (**This is a data file in Excel spreadsheet.**)

**DNA replication**

| Gene | $Log_2$ FC | *p*-vals |
|---|---|---|
| *CCNA2* | -0.821 | 9.37E-08 |
| *CDC45* | -0.533 | 9.64E-03 |
| *CDC6* | -0.553 | 6.34E-04 |
| *CDK2* | -0.600 | 6.88E-08 |
| *CDT1* | -1.065 | 1.28E-10 |
| *DBF4* | -0.630 | 5.79E-06 |
| *DNA2* | -0.602 | 2.65E-03 |
| *E2F1* | -0.522 | 3.06E-03 |
| *FEN1* | -0.763 | 7.24E-09 |
| *GINS1* | -0.623 | 1.88E-05 |
| *MCM10* | -0.795 | 3.15E-07 |
| *MCM2* | -0.736 | 2.98E-08 |
| *MCM3* | -0.644 | 1.69E-06 |
| *MCM4* | -0.698 | 8.50E-12 |
| *MCM5* | -0.836 | 3.48E-07 |
| *MCM6* | -0.635 | 7.73E-06 |
| *MCM7* | -0.929 | 5.76E-22 |
| *MCM8* | -0.631 | 1.22E-05 |
| *ORC6* | -0.502 | 4.38E-03 |
| *POLA2* | -0.541 | 6.50E-03 |
| *POLD3* | -0.545 | 3.37E-04 |
| *POLE* | -0.542 | 3.89E-03 |
| *PRIM1* | -0.654 | 1.06E-04 |
| *RFC5* | -0.539 | 7.35E-04 |
| *RPA3* | -0.663 | 5.82E-07 |
| *UBE2C* | -0.729 | 1.11E-12 |

**Transcription regulation mediated by p53**

| Gene | $Log_2$ FC | *p*-vals |
|---|---|---|
| *AURKA* | -0.756 | 2.06E-16 |
| *AURKB* | -0.822 | 2.69E-12 |
| *BBC3* | 0.594 | 5.11E-09 |
| *BIRC5* | -0.775 | 3.42E-15 |
| *BLM* | -0.562 | 1.85E-03 |
| *BRCA1* | -0.699 | 2.18E-07 |
| *BRIP1* | -0.640 | 7.24E-04 |
| *BTG2* | 0.789 | 5.61E-05 |
| *CCNA2* | -0.821 | 9.37E-08 |
| *CCNB1* | -0.563 | 1.96E-07 |
| *CDC25C* | -0.968 | 3.90E-08 |
| *CDK1* | -0.618 | 2.46E-09 |
| *CDK2* | -0.600 | 6.88E-08 |
| *CDKN1A* | 1.301 | 0.00E+00 |
| *DNA2* | -0.602 | 2.65E-03 |
| *E2F1* | -0.522 | 3.06E-03 |
| *EXO1* | -0.518 | 9.79E-03 |
| *FANCD2* | -0.633 | 1.18E-05 |
| *FANCI* | -0.677 | 2.68E-07 |
| *FOS* | -0.662 | 7.48E-06 |
| *GADD45A* | 1.134 | 6.23E-20 |
| *JUN* | 0.792 | 7.49E-13 |
| *MDC1* | -0.510 | 7.26E-03 |
| *MDM2* | 1.107 | 1.01E-35 |
| *PLK2* | 0.631 | 1.81E-07 |
| *PLK3* | 0.728 | 4.15E-04 |
| *RFC5* | -0.539 | 7.35E-04 |
| *RMI2* | -0.737 | 4.07E-05 |
| *RRAGD* | 0.562 | 4.96E-03 |
| *RRM2B* | 0.704 | 1.59E-06 |
| *SESN1* | 0.586 | 1.44E-03 |
| *SESN2* | 0.720 | 1.19E-07 |
| *TIGAR* | 0.863 | 2.45E-08 |
| *TP53I3* | 0.519 | 8.17E-05 |
| *TP53INP1* | 0.669 | 8.40E-04 |
| *TPX2* | -0.739 | 3.79E-19 |

**Table S6**

Three main gene clusters regulated by BTNL9 as identified by RNA-seq analysis.

**Table S6** (continue).

**Cell cycle**

| Gene | $Log_2$ FC | *p*-vals | Gene | $Log_2$ FC | *p*-vals | Gene | $Log_2$ FC | *p*-vals |
|---|---|---|---|---|---|---|---|---|
| *AAAS* | -0.544 | 5.94E-04 | *DHFR* | -0.722 | 8.46E-05 | *NCAPG* | -0.743 | 1.92E-07 |
| *AURKA* | -0.756 | 2.06E-16 | *DNA2* | -0.602 | 2.65E-03 | *NCAPH* | -0.731 | 4.55E-06 |
| *AURKB* | -0.822 | 2.69E-12 | *DSN1* | -0.603 | 6.84E-06 | *NCAPD3* | -0.650 | 6.38E-04 |
| *BIRC5* | -0.775 | 3.42E-15 | *E2F1* | -0.522 | 3.06E-03 | *NCAPG2* | -0.705 | 4.95E-07 |
| *BLM* | -0.562 | 1.85E-03 | *ESCO2* | -0.947 | 1.23E-09 | *NUF2* | -0.913 | 1.05E-12 |
| *BRIP1* | -0.640 | 7.24E-04 | *EXO1* | -0.518 | 9.79E-03 | *OIP5* | -0.929 | 5.39E-08 |
| *BRCA1* | -0.699 | 2.18E-07 | *FBXO5* | -0.737 | 1.66E-05 | *ORC6* | -0.502 | 4.38E-03 |
| *BRCA2* | -0.561 | 3.00E-04 | *FEN1* | -0.763 | 7.24E-09 | *PTTG1* | -0.865 | 3.27E-28 |
| *BUB1* | -0.797 | 1.24E-08 | *FOXM1* | -0.727 | 2.81E-05 | *PLK1* | -0.846 | 3.77E-18 |
| *BUB1B* | -0.525 | 2.06E-04 | *GTSE1* | -0.607 | 7.47E-05 | *POLA2* | -0.541 | 6.50E-03 |
| *CASC5* | -0.652 | 2.09E-05 | *GINS1* | -0.623 | 1.88E-05 | *POLE* | -0.542 | 3.89E-03 |
| *CKS1B* | -0.837 | 2.89E-27 | *H2AFV* | -0.549 | 2.94E-15 | *POLD3* | -0.545 | 3.37E-04 |
| *CDC20* | -0.545 | 9.99E-05 | *H2AFZ* | -0.752 | 2.93E-35 | *PRIM1* | -0.654 | 1.06E-04 |
| *CDC25B* | -0.536 | 4.96E-10 | *HIST1H2BK* | 0.582 | 6.94E-06 | *PRKAR2B* | -0.502 | 1.46E-02 |
| *CDC25C* | -0.968 | 3.90E-08 | *HIST1H4C* | -0.800 | 1.40E-44 | *PKMYT1* | -0.786 | 1.30E-05 |
| *CDC45* | -0.533 | 9.64E-03 | *HIST2H2AC* | -0.529 | 5.20E-03 | *RAD21* | -0.581 | 7.42E-15 |
| *CDC6* | -0.553 | 6.34E-04 | *HJURP* | -0.799 | 6.45E-16 | *RHNO1* | -0.540 | 9.91E-04 |
| *CDCA5* | -0.771 | 5.11E-05 | *HMMR* | -0.777 | 1.47E-12 | *RMI2* | -0.737 | 4.07E-05 |
| *CDCA8* | -0.824 | 6.63E-09 | *ITGB3BP* | -0.931 | 5.78E-13 | *RFC5* | -0.539 | 7.35E-04 |
| *CENPA* | -0.874 | 2.34E-06 | *KIF18A* | -0.625 | 4.82E-04 | *RPA3* | -0.663 | 5.82E-07 |
| *CENPE* | -0.800 | 2.77E-17 | *KIF20A* | -0.826 | 7.39E-08 | *RRM2* | -0.734 | 3.26E-13 |
| *CENPF* | -0.868 | 0.00E+00 | *KIF23* | -0.835 | 2.20E-11 | *SUN2* | -0.502 | 3.15E-03 |
| *CENPH* | -0.700 | 1.62E-04 | *KIF2C* | -0.782 | 6.48E-07 | *SGOL1* | -0.713 | 2.31E-06 |
| *CENPK* | -0.755 | 2.49E-06 | *KNTC1* | -0.728 | 8.40E-05 | *SGOL2* | -0.699 | 3.41E-09 |
| *CENPM* | -0.598 | 1.71E-03 | *LMNB1* | -0.693 | 1.83E-09 | *SPC25* | -0.577 | 1.53E-03 |
| *CENPN* | -0.727 | 6.14E-07 | *MAD2L1* | -0.718 | 7.07E-05 | *SKA1* | -0.615 | 1.86E-03 |
| *CENPU* | -0.525 | 2.74E-03 | *MDM2* | 1.107 | 1.01E-35 | *SMC2* | -0.781 | 2.05E-17 |
| *CENPW* | -0.568 | 8.61E-04 | *MDC1* | -0.510 | 7.26E-03 | *SMC4* | -0.680 | 9.51E-29 |
| *CEP152* | -0.535 | 1.20E-04 | *MCM10* | -0.795 | 3.15E-07 | *TK1* | -1.089 | 1.95E-12 |
| *CDT1* | -1.065 | 1.28E-10 | *MCM8* | -0.631 | 1.22E-05 | *TMPO* | -0.556 | 6.64E-10 |
| *CLSPN* | -0.771 | 8.76E-08 | *MCM2* | -0.736 | 2.98E-08 | *TOP2A* | -0.867 | 0.00E+00 |
| *CCNA2* | -0.821 | 9.37E-08 | *MCM3* | -0.644 | 1.69E-06 | *TPX2* | -0.739 | 3.79E-19 |
| *CCNB1* | -0.563 | 1.96E-07 | *MCM4* | -0.698 | 8.50E-12 | *TUBA1A* | 0.735 | 2.32E-08 |
| *CCNB2* | -0.967 | 2.91E-15 | *MCM5* | -0.836 | 3.48E-07 | *TUBA1B* | -0.510 | 5.00E-25 |
| *CCND1* | 0.629 | 1.25E-16 | *MCM6* | -0.635 | 7.73E-06 | *UBE2C* | -0.729 | 1.11E-12 |
| *CDK1* | -0.618 | 2.46E-09 | *MCM7* | -0.929 | 5.76E-22 | *MYBL2* | -0.550 | 8.00E-06 |
| *CDK2* | -0.600 | 6.88E-08 | *MIS18BP1* | -0.771 | 1.10E-11 | *VRK1* | -0.502 | 1.32E-03 |
| *CDKN1A* | 1.301 | 0.00E+00 | *NDC80* | -0.596 | 1.61E-03 | *WHSC1* | -0.617 | 1.91E-09 |
| *CDKN2C* | -0.883 | 2.14E-06 | *NEK2* | -0.738 | 1.17E-06 | *ZWINT* | -0.713 | 1.53E-06 |
| *DBF4* | -0.630 | 5.79E-06 | *NCAPD2* | -0.699 | 2.40E-12 | | | |

**Table S7**

Overlapping genes between 9,707 genes from ChIP-seq analysis and 444 DEGs from RNA-seq in A549 cells. (**This is a data file in Excel spreadsheet.**)

**RT-qPCR**

| Primer name | Sequence (5' to 3') | Amplicon size (bp) |
|---|---|---|
| AURKA-F | cttcttggatcagctggagagc | 170 |
| AURKA-R | caaagaactccaaggctccaga | |
| AURKB-F | agtgggacacccgacatcttaa | 158 |
| AURKB-R | tctatctgggacttgaagaggacc | |
| BBC3-F | ACGACCTCAACGCACAGTACGA | 147 |
| BBC3-R | CCTAATTGGGCTCCATCTCGGG | |
| BTNL9-F | CCATTCAGAACCTGCTGCTGAGC | 114 |
| BTNL9-R | GCAGTGGCAGTGTAGCAACAAAGG | |
| CCNA2-F | gaatatcaacccggaaaaggcag | 171 |
| CCNA2-R | ggaacggtgacatgctcatcat | |
| CCNB1-F | caaagtcagtgaacaactgcagg | 174 |
| CCNB1-R | tcaggttctggctcaggttct | |
| CDC25C-F | AGAAGCCCATCGTCCCTTTGGA | 133 |
| CDC25C-R | GCAGGATACTGGTTCAGAGACC | |
| CDK1-F | ggaaaccaggaagcctagcatc | 124 |
| CDK1-R | ggatgattcagtgccattttgcc | |
| CDK2-F | cggagttgtgtacaaagccaga | 173 |
| CDK2-R | ctgtgtgaatgacatccagcagc | |
| E2F1-F | GGACCTGGAAACTGACCATCAG | 121 |
| E2F1-R | CAGTGAGGTCTCATAGCGTGAC | |
| FOXM1-F | TCTGCCAATGGCAAGGTCTCCT | 145 |
| FOXM1-R | CTGGATTCGGTCGTTTCTGCTG | |
| GADD45A-F | CTGGAGGAAGTGCTCAGCAAAG | 146 |
| GADD45A-R | AGAGCCACATCTCTGTCGTCGT | |
| MCM2-F | TGCCAGCATTGCTCCTTCCATC | 163 |
| MCM2-R | AAACTGCGACTTCGCTGTGCCA | |
| MCM3-F | CGAGACCTAGAAAATGGCAGCC | 107 |
| MCM3-R | GCAGTGCAAAGCACATACCGCA | |
| MCM7-F | GCCAAGTCTCAGCTCCTGTCAT | 136 |
| MCM7-R | CCTCTAAGGTCAGTTCTCCACTC | |
| MDM2-F | TGTTTGGCGTGCCAAGCTTCTC | 113 |
| MDM2-R | CACAGATGTACCTGAGTCCGATG | |
| MYBL2-F | CACCAGAAACGAGCCTGCCTTA | 130 |
| MYBL2-R | CTCAGGTCACACCAAGCATCAG | |
| ORC6-F | ccagacacagcaagtggatcttg | 125 |
| ORC6-R | ttacaccggatgtggctaccat | |
| GAPDH-F | ACATCGCTCAGACACCATG | 143 |
| GAPDH-R | TGTAGTTGAGGTCAATGAAGGG | |

**Table S8**
Primers list.

**Table S8** (continue).

**ChIP-qPCR**

| Primer name | Sequence (5’ to 3)' | Amplicon size |
|---|---|---|
| DDX1-F | cttccatcggaaccgttctcc | 186 |
| DDX1-R | cagagtttccacaaacgcaccg | |
| KANSL3-F | cgtgttcttggctctcctcttc | 117 |
| KANSL3-R | caccgagtaaaacgcggaagc | |
| UBC-F | ggatttgggtcgcagttcttgttt | 174 |
| UBC-R | cagttcagggcaaccttgctc | |

**Plasmid construction**

| Primer name | Sequence (5' to 3') | Amplicon size |
|---|---|---|
| BTNL9-AMtag-F | GTCAGATCCGCTAGCGTCGCCACCATGGTCGATCTGAG | 1782 |
| BTNL9-AMtag-R | TCGAGGTCGAGAATTCTAGTAAGCCTGAGAGAGTATCACG | |
| BTNL9-WT-F | GTCAGATCCGCTAGCGTCGCCACCATGGTCGAT | 1664 |
| BTNL9-WT-R | TGTCTCGAGGTCGAGAATTCTACCACCAGTCCAGAGCAGGATCA | |

| Antibody | Cat number | Source | Identifier |
| --- | --- | --- | --- |
| Am-Tag | 91111 | Active Motif | RRID:AB_2793779 |
| BBC3 | 55120-1-AP | Proteintech | RRID:AB_10859944 |
| BTNL9 | LS-C399149 | LS-Bio | |
| CASP9 | 9502S | Cell Signaling | RRID:AB_2068621 |
| CDC25C | 16485-1-AP | Proteintech | RRID:AB_2291330 |
| CDK1 | 19532-1-AP | Proteintech | RRID:AB_10638617 |
| CDK2 | A0094 | Abclonal | RRID:AB_2861449 |
| CCNA2 | A19036 | Abclonal | RRID:AB_2862528 |
| CCNB1 | A19037 | Abclonal | RRID:AB_2862529 |
| CDKN1A | 2947T | Cell Signaling | RRID:AB_823586 |
| E2F1 | 32-1400 | Thermo Fisher Scientific | RRID:AB_2533065 |
| FOXM1 | 13147-1-AP | Proteintech | RRID:AB_2106213 |
| GADD45A | A11768 | Abclonal | RRID:AB_2758745 |
| H3K27ac | MA5-23516 | Thermo Fisher Scientific | RRID:AB_2608307 |
| MCM2 | 10513-1-AP | Proteintech | RRID:AB_2142131 |
| MCM3 | 15597-1-AP | Proteintech | RRID:AB_2141973 |
| MCM7 | 11225-1-AP | Proteintech | RRID:AB_2297584 |
| MDM2 | 66511-1-Ig | Proteintech | RRID:AB_2881874 |
| MYBL2 | 18896-1-AP | Proteintech | RRID:AB_10644283 |
| ORC6 | 17784-1-AP | Proteintech | RRID:AB_2283320 |
| TP53 | 2527 | Cell Signaling | RRID:AB_10695803 |
| YY1 | 63227 | Cell Signaling | RRID:AB_2799641 |
| β-Actin | 4970 | Cell Signaling | RRID:AB_2223172 |
| Goat anti-mouse HRP | 115-035-003 | Jackson Immunoresearch | RRID:AB_10015289 |
| Goat anti-rabbit HRP | 111-035-003 | Jackson Immunoresearch | RRID:AB_2313567 |
| Goat anti-rabbit IgG Alexa Fluor® 594-conjugated | A32740 | Thermo Fisher Scientific | RRID:AB_2762824 |
| Mouse IgG2a Isotype Control | 02-6200 | Thermo Fisher Scientific | RRID:AB_2532943 |

**Table S9**
Antibodies list.
